# SATAY-Based Chemogenomic Screening uncovers Antifungal Resistance Mechanisms and Key Determinants of ATI-2307 and Chitosan Sensitivity

**DOI:** 10.1101/2024.09.20.614104

**Authors:** Matthew T. Karadzas, Agnès H. Michel, Andreas Mosbach, George Giannakopoulos, Ruairi McGettigan, Gabriel Scalliet, Benoît Kornmann

## Abstract

Multidrug-resistant fungal pathogens are a serious threat to public health and global food security. Mitigation requires the discovery of antifungal compounds with novel modes of action, along with a comprehensive understanding of the molecular mechanisms governing antifungal resistance. Here, we apply SAturated Transposon Analysis in Yeast (SATAY), a powerful transposon sequencing method in *Saccharomyces cerevisiae*, to uncover loss- and gain-of-function mutations conferring resistance to 20 different antifungal compounds. These screens identify a diverse array of novel resistance mechanisms and multiple modes of action. SATAY is performed in drug-sensitive strains to generate chemogenomic profiles for compounds that lack activity against conventional laboratory strains. This study therefore provides a significant resource for exploring cellular responses to chemical stresses. We discover that the natural antifungal Chitosan electrostatically interacts with cell wall mannosylphosphate, and that the transporter Hol1 concentrates the novel antifungal ATI-2307 within yeast. This latter finding presents an avenue for drug design initiatives, but also unveils a straightforward evolutionary path to ATI-2307 resistance with minimal fitness cost.

## Introduction

Fungal pathogens are a serious public health concern. Invasive fungal infections (IFIs) cause approximately 2.5 million deaths worldwide every year (Denning, 2024), and cases of IFIs are continuing to rise sharply, concomitant with the expanding population of immunocompromised individuals (Bongomin et al., 2017). Meanwhile, fungal plant pathogens pose the most serious biotic challenge to global food security (Steinberg & Gurr, 2020), accounting for approximately 20% of crop yield losses pre-harvest, and a further 10-20% post-harvest (Fisher et al., 2018). Fungal pathogens are currently controlled using a very limited number of antifungal compounds: only four classes of clinical antifungals are available (Puumala et al., 2024), whilst the agrochemical fungicide market comprises of six main classes (Fones et al., 2020). Resistance to all these classes has emerged in major fungal pathogens, undermining our ability to control fungal infections in both clinical and agricultural settings (Fisher et al., 2018; Perlin et al., 2017). Meanwhile, our understanding of the molecular mechanisms responsible for antifungal resistance remains incomplete, yet this knowledge is critical for the development of effective resistance management strategies (Robbins et al., 2017). Furthermore, the scarcity of antifungal compounds demands concerted action towards the discovery of antifungals with novel modes of action (Perfect, 2017; Puumala et al., 2024).

Chemogenomic screening approaches in the model yeast *Saccharomyces cerevisiae* have accelerated the discovery of antifungal resistance mechanisms and modes of action (Giaever et al., 1999, 2004; Hoepfner et al., 2014; Lee et al., 2014). These approaches involve the systematic identification of every gene that promotes antifungal sensitivity or resistance, and are enabled by the fact that, unlike fungal pathogens, *S. cerevisiae* is easy to genetically manipulate, has an extensively annotated genome, a complete sexual cycle and grows rapidly on cheap medium under standard laboratory conditions (Demuyser & Van Dijck, 2019; Duina et al., 2014). Genome-wide yeast deletion collections (Giaever et al., 2002; Winzeler et al., 1999) have been successfully utilised for chemogenomic screening approaches such as haploinsufficiency profiling (HIP) and homozygous profiling (HOP) (Giaever et al., 1999, 2004; Hillenmeyer et al., 2008; Hoepfner et al., 2014; Parsons et al., 2004). However, yeast deletion collections are only available in a small number of conventional laboratory strains, and since genetic background affects the penetrance and/or expression of genetic traits, this can complicate the connection from genotype to phenotype. These analyses are complicated further by the fact that many deletion collection strains have accumulated secondary mutations (Teng et al., 2013).

Recently, an innovative genome-wide screening approach, generically termed transposon sequencing (Tn-seq), has been developed in a variety of microorganisms, including *S. cerevisiae* (Gale et al., 2020; Gao et al., 2018; Guo et al., 2013; Li et al., 2011; Mielich et al., 2018; Patterson et al., 2018; Sanchez et al., 2019; Segal et al., 2018; van Opijnen et al., 2009; Zhu et al., 2018; Edskes et al., 2018; Michel et al., 2017). Tn-seq utilises random transposon mutagenesis to generate dense transposon libraries, in which every gene is disrupted by multiple independent transposon insertions. The insertion sites are identified *en masse* using next-generation sequencing, with the number of sequencing reads for each insertion reflecting the abundance of the corresponding mutants in the library population. When a library is grown under a test condition, the change in the number of reads for each insertion reveals the effect of the insertion on fitness (van Opijnen et al., 2009). In this way, transposon libraries treated with antimicrobial compounds can identify genes affecting drug susceptibility (Coe et al., 2019; Gallagher et al., 2011; Li et al., 2023; Murray et al., 2015). Indeed, recent studies have utilised Tn-seq in the fungal pathogen *Candida glabrata* to identify genes affecting susceptibility to Fluconazole (Gale et al., 2020; Gale et al., 2023), Micafungin (Nickels et al., 2024) and the calcineurin inhibitor FK506 (Pavesic et al., 2024). Tn-seq has also been used to explore the genetic basis of Fluconazole susceptibility in *Candida albicans* (Gao et al., 2018). Tn-seq overcomes some of the main drawbacks of yeast deletion collections: transposon libraries are easily generated *de novo*, facilitating genome-wide screening in different genetic backgrounds (Levitan et al., 2020; Michel et al., 2017), and secondary mutations arising during Tn-seq do not confound data interpretation, since each gene is interrupted by multiple independent insertions (Michel et al., 2017).

SAturated Transposon Analysis in Yeast (SATAY) is a powerful Tn-seq method in *S. cerevisiae* (Chen et al., 2022; Michel et al., 2017; Michel & Kornmann 2022, John Peter et al., 2022) with significant potential for chemogenomic screening. Indeed, SATAY can simultaneously assess the fitness effects of loss- and gain-of-function mutations in different strains and test conditions in a labour- and time-efficient manner (Michel et al., 2017; Serbyn et al., 2020). Moreover, SATAY can be easily multiplexed (John Peter et al., 2022; Michel & Kornmann, 2022).

To address gaps in our understanding of antifungal resistance and modes of action, which is critical for future fungal disease control, we apply SATAY for chemogenomic screening of 20 antifungal compounds. This selection includes major classes of clinical and agrochemical antifungals as well as experimental compounds, targeting a range of cellular processes, such as cell wall biosynthesis, ergosterol biosynthesis, protein synthesis, glycosylation and oxidative phosphorylation. The selection also includes compounds with unclear modes of action. Furthermore, by performing SATAY in drug-sensitive strains, we reveal the modes of action for compounds that lack activity against conventional laboratory strains, such as Fludioxonil, Iprodione and Chitosan. In addition, our screen with ATI-2307 unveils the genetic determinants underpinning the potency of this novel antifungal.

## Results

### 1. SATAY can identify loss- and gain-of-function mutations conferring antifungal sensitivity or resistance

To evaluate the effectiveness of SATAY for chemogenomic screening, we performed a series of proof-of-principle screens with well-characterised antifungal compounds. Transposon libraries (Supplementary Table 1) were grown for two successive rounds from OD_600_ 0.1-0.2 to saturation in the presence of sub-lethal concentrations of antifungal compounds inhibiting growth by about 30% (∼IC_30_) (Supplementary Table 2). This concentration optimised the likelihood of identifying transposon insertions conferring antifungal resistance or sensitivity in the same screen (Hoepfner et al., 2014).

Transposon insertions interrupting the coding DNA sequence (CDS) of genes revealed loss-of-function mutations conferring antifungal resistance. For example, Amphotericin B resistance was caused by transposons disrupting ergosterol homeostasis (**Figure 1A, orange**), the Target Of Rapamycin Complex 1 (TORC1) (**Figure 1A, pink**) and the TORC1-activating EGO complex (**Figure 1A, blue**), consistent with previous reports (Bojsen et al., 2016; Daum et al., 1999; Martel et al., 2010). Caspofungin resistance was conferred by transposons in sphingolipid biosynthesis genes (**Figure 1B, orange**) (Supplementary Table 3), aligning with its known mode of action (García et al., 2015; Healey et al., 2012; Satish et al., 2019). Caspofungin also positively selected transposons in *ERG3*, *LEM3*, *CRZ1*, and *GSC2* (**Figure 1B, blue and pink**), in agreement with previous findings (García et al., 2015; Lesage et al., 2004; Markovich et al., 2004). Myriocin, a potent inhibitor of serine palmitoyltransferase (Miyake et al., 1995), positively selected transposons in genes associated with cellular trafficking (**Figure 1C**: GARP complex, pink; retromer, teal; vesicular transport, orange), phospholipid translocation (**Figure 1C**, **blue**) and chromatin remodelling (**Figure 1C, brown**). Consistently, these genes have previously been associated with Myriocin susceptibility (Fröhlich et al., 2015).

**Figure 1.**
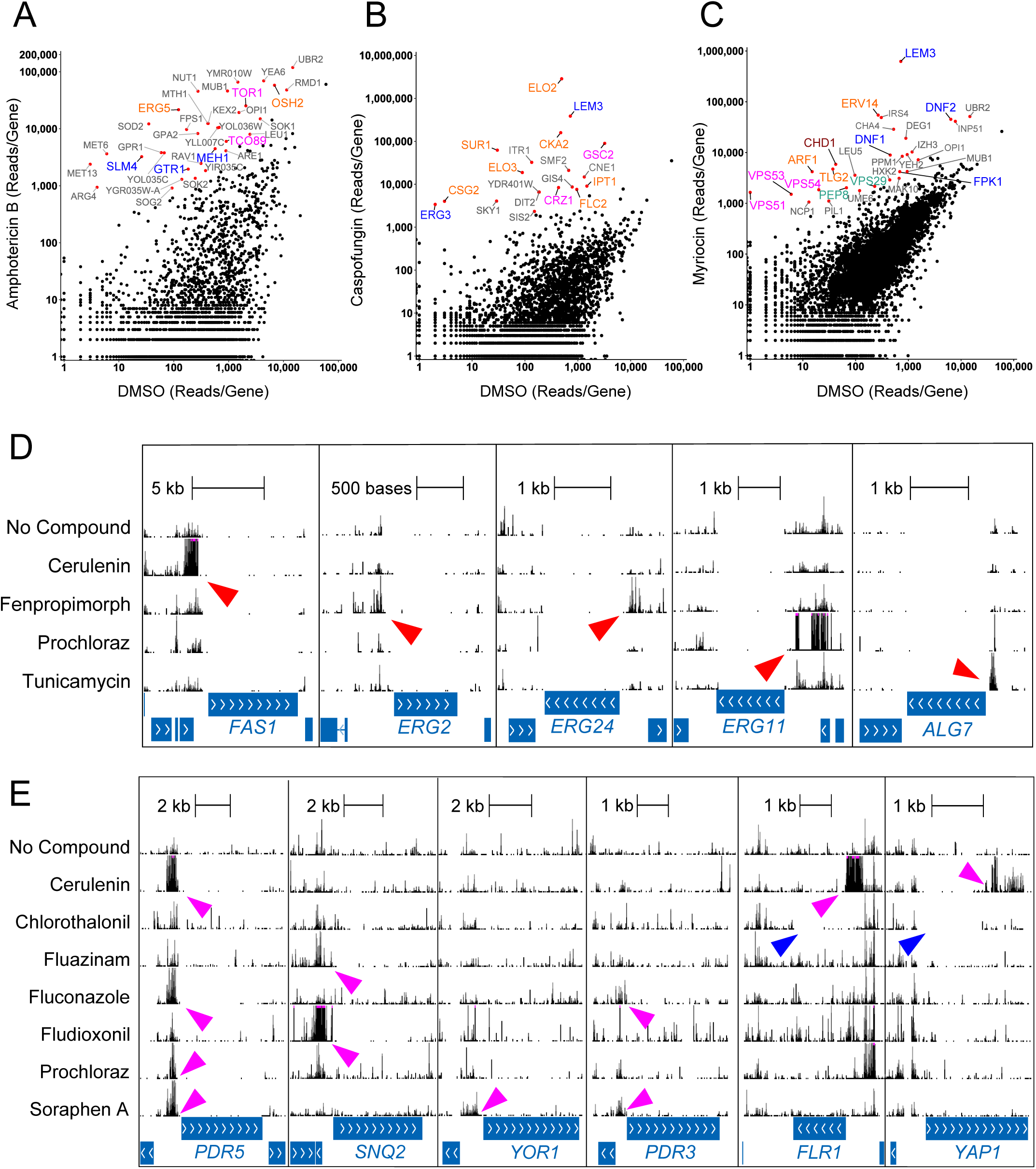
SATAY can identify loss- and gain-of-function mutations conferring antifungal sensitivity or resistance. **A-C)** Scatter plots comparing the number of sequencing reads mapping to each of the 6603 yeast coding DNA sequences (CDSs) in transposon libraries treated with DMSO versus **A)** Amphotericin B (0.375 μg/mL) **B)** Caspofungin (0.015 μg/mL) **C)** Myriocin (0.2 μg/mL). Each dot represents a single gene. Gene names are colour-coded by biological process. **A)** Orange - ergosterol homeostasis, pink - TORC1 complex subunits, blue – EGO complex subunits. **B)** Orange – sphingolipid biosynthesis, pink – β-1,3-glucan biosynthesis, blue – membrane lipid homeostasis. **C)** Orange – vesicular-mediated transport, pink – GARP complex subunits, blue – phospholipid translocation, teal – retromer subunits, maroon – chromatin remodelling. **D)** Transposon insertion maps for five antifungal drug targets. The height of the bars corresponds to the number of sequencing reads, shown on a linear scale that is capped at 500. Horizontal blue bars represent CDSs. Red arrowheads indicate positively selected insertions in the promoter regions of genes encoding drug targets. **E)** Transposon insertion maps for four drug efflux pumps (*PDR5*, *SNQ2*, *YOR1*, *FLR1*) and two transcriptional regulators of the PDR network (*PDR3*, *YAP1*). The height of the bars corresponds to the number of sequencing reads, shown on a linear scale that is capped at 500. Pink arrowheads indicate positively selected insertions in promoter regions, or in the *YAP1* inhibitory domain. Blue arrowheads indicate regions of *FLR1* and *YAP1* where insertions were negatively selected.

Antifungal resistance can also be conferred by gain-of-function mutations leading to target overexpression (Robbins et al., 2017). Given that the SATAY transposon bears cryptic promoter sequences (Yu et al., 2004), insertions in promoters can lead to downstream gene overexpression (Serbyn et al., 2020). We therefore wondered whether antifungal compounds might positively select transposons in the promoters of target genes. Since the annotation of promoters in *S. cerevisiae* is incomplete, we performed three separate analyses on insertions mapping within 1) 200-bp upstream of the ATG start codon 2) 500-bp upstream of the ATG start codon and 3) 200-bp upstream of the transcription start site, where known (Xu et al., 2009). For transposon libraries separately treated with Cerulenin, Fenpropimorph, Prochloraz or Tunicamycin, the promoter regions containing the most enriched transposons were precisely those upstream of the genes encoding the corresponding targets: *FAS1* for Cerulenin (Johansson et al., 2008), *ERG2* and *ERG24* for Fenpropimorph (Marcireau et al., 1990), *ERG11* for Prochloraz (Becher & Wirsel, 2012) and *ALG7* for Tunicamycin (Tkacz & Lampen, 1975) (**Figure 1D, red arrowheads**) (Supplementary Figures S3-5). This demonstrates that SATAY can be used to identify the direct targets of antifungal compounds.

In addition, our analysis of promoter regions revealed that SATAY can identify the drug efflux pumps responsible for detoxifying antifungal compounds (Supplementary Figure S3-5). The overexpression of ATP-binding cassette (ABC) (Prasad & Goffeau, 2012) and Major Facilitator Superfamily (MFS) transporters (Sá-Correia et al., 2009) is the basis for pleiotropic drug resistance (PDR) in *S. cerevisiae* (Gulshan & Moye-Rowley, 2007). We observed that transposons in the promoter of *PDR5*, encoding an ABC transporter (Meyers et al., 1992), were selected by Cerulenin, Fluconazole, Prochloraz and Soraphen A (**Figure 1E, pink arrowheads**), all known Pdr5 substrates (Arya et al., 2019; Kolaczkowski et al., 1998; Rogers et al., 2001). Fluconazole and Soraphen A also positively selected transposons in the promoter of *PDR3*, a transcription factor that regulates *PDR5* expression (Moye-Rowley, 2003; Sá-Correia et al., 2009). Similarly, *SNQ2* promoter insertions were selected by Fluazinam and Fludioxonil (**Figure 1E, pink arrowheads**), suggesting that this ABC transporter (Servos et al., 1993) exports these compounds. Notably, the positive selection of transposons in the promoters of *PDR5* and *SNQ2* correlated with the negative selection of transposons in their CDSs.

Interestingly, SATAY reveals insights into the regulation of drug efflux pumps, which are only expressed under specific stress conditions. The loss of non-expressed pumps has no effect on fitness but, during drug treatments, their increased expression becomes beneficial, leading to positive selection of transposons in their promoters. This was evident for *FLR1*, which encodes an MFS transporter, after Cerulenin treatment (**Figure 1E, pink arrowheads**). Conversely, for efflux pumps that are sufficiently expressed, insertions in the promoter have little consequence, but insertions in the CDS are detrimental. This is the case for *FLR1* during Chlorothalonil treatment, where insertions in the promoter are neutral, but insertions in the CDS are negatively selected (**Figure 1E, blue arrowhead**). We therefore conclude that *FLR1* expression is silent upon Cerulenin treatment, but strong in Chlorothalonil. *FLR1* is transcriptionally regulated by the redox-sensitive transcription factor Yap1 (Alarco et al., 1997; Jungwirth et al., 2000). Yap1 contains a C-terminal inhibitory domain that is inactivated by oxidative stress (Gulshan et al., 2005; Kuge et al., 1997). Accordingly, Cerulenin favours insertions that truncate Yap1’s C-terminus (**Figure 1E, pink arrowhead**), leading to its constitutive activation and subsequent *FLR1* overexpression (Alarco et al., 1997). Meanwhile, Chlorothalonil likely activates Yap1 by inducing an oxidative stress (Tillman et al., 1973). Supporting this, insertions in the N-terminal transcription activator region of *YAP1* CDS are negatively selected on Chlorothalonil, while insertions in the inhibitory domain show no significant selection (**Figure 1E**, **blue arrowhead**). Therefore, the contrasting insertion profiles for *FLR1* and *YAP1* on Cerulenin and Chlorothalonil reveal the ability of Yap1 to sense Chlorothalonil and its inability to sense Cerulenin.

In summary, our proof-of-principle screens validate that SATAY is a robust approach for chemogenomic screening. We have demonstrated that SATAY can reveal loss- and gain-of-function mutations conferring antifungal resistance, the direct targets of antifungal compounds and the efflux pumps mediating drug export. Genes conferring resistance or sensitivity to all compounds screened in this study are shown in Supplementary Figures S1-5. Genomic maps of the transposon insertions can be browsed here: https://genome.ucsc.edu/s/Matthew%20Karadzas/Histograms_Karadzas%202024 https://genome.ucsc.edu/s/Matthew%20Karadzas/Transposon_Maps_Karadzas_2024

### 2. SATAY can be performed in drug-sensitive strains of *S. cerevisiae*

Although *S. cerevisiae* is a powerful model for chemogenomic screening (Robbins & Cowen, 2022), it lacks sensitivity to several antifungal compounds. One reason for this is the PDR network (Kolaczkowski et al., 1998), which can mask cellular responses relevant to a compound’s mode of action and demands high levels of compound use. This is illustrated by our wild-type (ByK352) transposon library treated with 10 µg/mL Prochloraz, an imidazole fungicide that is a substrate of the PDR network (Kolaczkowski et al., 1998). Indeed, most of the top-hits in this screen were associated with the Gene Ontology (GO) terms “mitochondrial gene expression” and “mitochondrial translation” (**Figure 2A)** (Supplementary Table 3) because mutations disrupting these functions activate the PDR network (Hallstrom & Moye-Rowley, 2000; Zhang & Moye-Rowley, 2001), increasing Prochloraz detoxification.

**Figure 2.**
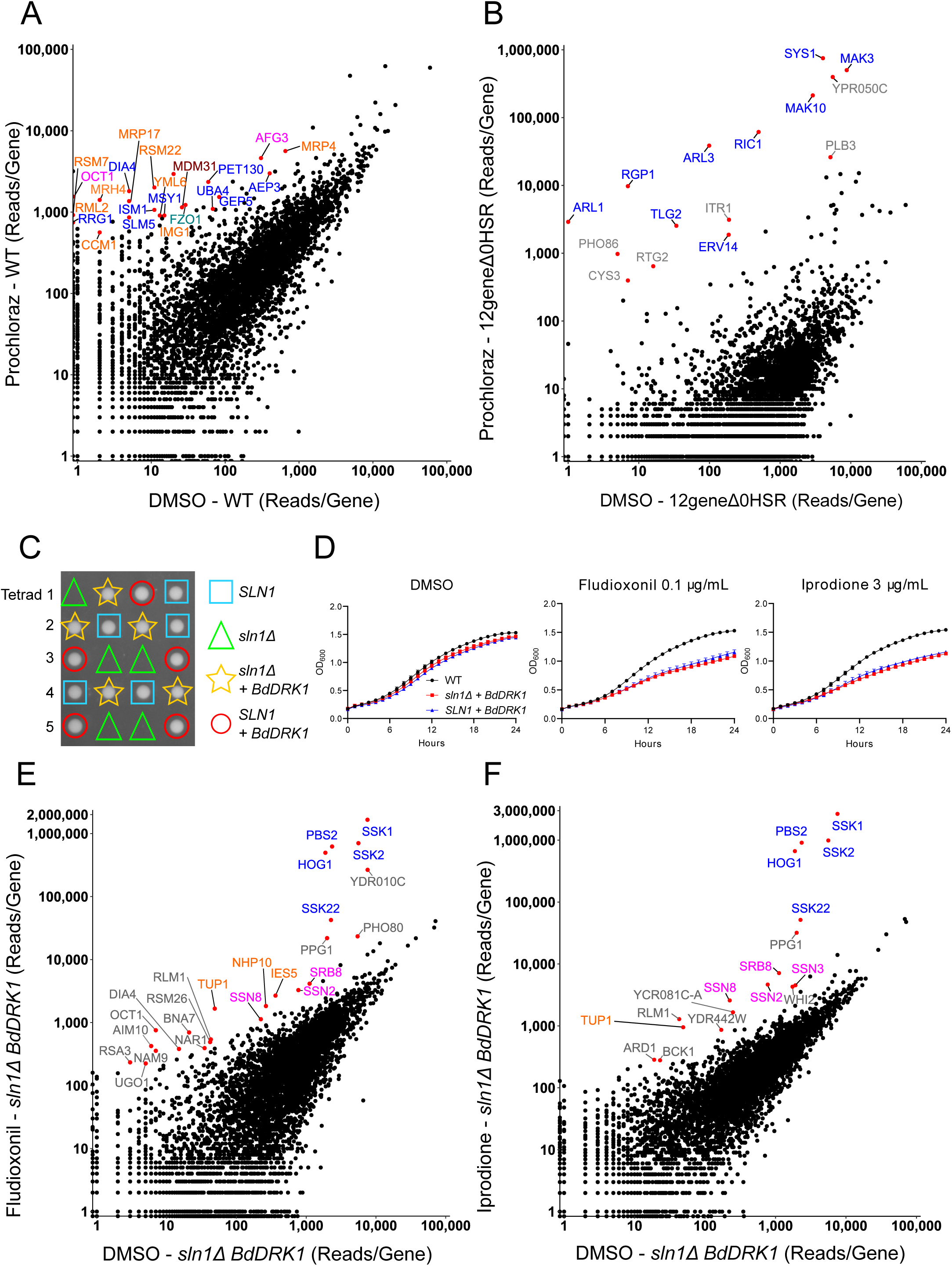
SATAY can be performed in drug-sensitive strains of *S. cerevisiae*. **A)** Scatter plot comparing the number of sequencing reads mapping to each of the 6603 yeast CDSs in a PDR-proficient transposon library treated with DMSO versus Prochloraz (10 μg/mL). Each dot represents a single gene. Gene names are colour-coded by biological process: orange – mitochondrial gene expression, blue – mitochondrial translation, pink – mitochondrial protein processing, teal – mitochondrial genome maintenance, maroon – mitochondrial phospholipid homeostasis. **B)** As in **(A)**, for a PDR-attenuated (12geneΔ0HSR) transposon library treated with DMSO versus Prochloraz (0.4 μg/mL). Genes involved with vesicle-mediated transport are coloured blue. **C)** Tetrad dissection of *SLN1/sln1*Δ *BdDRK1*+/- heterozygote on YPD medium. The genotypes of the non-growing spores have been inferred from the genotypes of the growing ones, assuming a Mendelian meiotic segregation pattern. **D)** Growth curves of wild-type, *sln1*Δ *BdDRK1* and *SLN1 BdDRK1* strains in SC 2% glucose medium containing either 1% DMSO (v/v), Fludioxonil (0.1 μg/mL) or Iprodione (3 μg/mL). OD_600_ measurements were recorded every 20 minutes using a Bioscreen C™ instrument. Each data point is the mean OD_600_ measurement for three independent biological replicates. Error bars show standard error of mean (SEM). **E-F)** Scatter plots comparing the number of sequencing reads mapping to each of the 6603 yeast CDSs in a *sln1*Δ *BdDRK1* transposon library treated with **(E)** Fludioxonil (0.1 μg/mL) or **(F)** Iprodione (3 μg/mL). Gene names are colour-coded by biological process: blue – HOG signalling pathway, pink – RNA polymerase holoenzyme or mediator complex subunit, orange – chromatin remodelling. Note that *YDR010C* is a dubious open reading frame (ORF) located in the promoter of the drug-efflux pump *SNQ2* (Figure 1E).

We therefore assessed whether SATAY can be performed in a PDR-attenuated strain called 12geneΔ0HSR, in which eight ABC transporters and four transcriptional regulators of the PDR network have been deleted (Chinen et al., 2011). Consistently, 12geneΔ0HSR was 25-fold more sensitive towards Prochloraz than the wild-type strain (ByK352) also derived from BY4741 (Supplementary Table 2). Compared to the ByK352 transposon library, Prochloraz positively selected transposons in an entirely different set of genes in the 12geneΔ0HSR transposon library (**Figure 2B**). Indeed, many of the top-hits were associated with the GO term “vesicle-mediated transport” (Supplementary Table 3). In fact, five of the most enriched genes (*MAK3*, *MAK10*, *ARL3*, *SYS1, ARL1*) all act in the same signalling pathway that culminates in the Golgi recruitment of Arl1, which promotes selective vesicle trafficking at the trans-Golgi network (TGN) (Behnia et al., 2004; Setty et al., 2003; Yu & Lee, 2017). Similarly, a recent Tn-seq study in a PDR-attenuated strain of *Candida glabrata* also observed that transposon insertions in genes with roles at the Golgi, including *MAK3* and *MAK10*, conferred Fluconazole resistance (Gale et al., 2023). Thus, SATAY can be performed in PDR-attenuated strains to unveil PDR-independent resistance mechanisms at lower drug concentrations.

Another reason why *S. cerevisiae* lacks sensitivity to several compounds is that it does not express the corresponding target. Sensitivity can often be engineered by heterologously expressing the target in *S. cerevisiae*, and since SATAY can be performed in genetically engineered strains, it is a promising approach for the chemogenomic analysis of these compounds (John Peter et al., 2022; Michel et al., 2017). The fungicides Fludioxonil and Iprodione lack activity against *S. cerevisiae* (Kilani & Fillinger, 2016; Leroux et al., 1992; Motoyama et al., 2005). In filamentous fungi, these compounds are sensed by group III hybrid histidine kinases (HHKs), leading to the constitutive activation of the High Osmolarity Glycerol (HOG) signalling pathway, which is lethal (Fillinger et al., 2012; Furukawa et al., 2012; Motoyama et al., 2005; Kojima et al., 2004; Zhang et al., 2002). There are conflicting ideas regarding how Fludioxonil affects group III HHKs, and a recent model proposes that Fludioxonil does not target group III HHKs directly, but via an upstream factor of unclear identity (Lawry et al., 2017). *S. cerevisiae* does not harbour group III HHKs, but encodes one essential group VI HHK called Sln1 (Catlett et al., 2003; Maeda et al., 1994; Ota & Varshavsky, 1993; Posas et al., 1996). *S. cerevisiae* strains heterologously expressing a group III HHK are sensitive to Fludioxonil and Iprodione (Buschart et al., 2012; Furukawa et al., 2012; Lawry et al., 2017; Meena et al., 2010; Motoyama et al., 2005), indicating that if Fludioxonil indeed acts upon an upstream factor, it must be present in *S. cerevisiae*, and SATAY might therefore be an appropriate way to identify it. We expressed a group III HHK called Drk1 from the fungal pathogen *Blastomyces dermatitidis* (Nemecek et al., 2006) in *S. cerevisiae*, which complemented the otherwise lethal deletion of *SLN1* (**Figure 2C**). We found that *sln1*Δ *BdDRK1* strains are equally susceptible to Fludioxonil and Iprodione as *SLN1 BdDRK1* strains (**Figure 2D**), consistent with reports that Fludioxonil activates the HOG pathway in a dominant manner (Lawry et al., 2017; Randhawa et al., 2019). We generated a transposon library in a *sln1*Δ *BdDRK1* strain, in which the absence of Sln1 ensured that Fludioxonil and Iprodione resistance could not result from loss of *BdDRK1*.

Fludioxonil and Iprodione strongly selected insertions in the five non-essential components of the HOG pathway (*SSK1*, *SSK2*, *SSK22*, *PBS2* and *HOG1*) in the *sln1*Δ *BdDRK1* transposon library (**Figure 2E-F, blue**)(Hohmann, 2002). This aligns with studies showing that mutations in HOG pathway components confer Fludioxonil and Iprodione resistance (Buschart et al., 2012; Furukawa et al., 2012; Kojima et al., 2004; Motoyama et al., 2005; Zhang et al., 2002). We also observed that Fludioxonil and Iprodione selected transposons in subunits of the RNA polymerase II holoenzyme (*SSN8*) and mediator complex (*SSN2*, *SRB8*), subunits of the INO80 chromatin remodelling complex (*IES5*, *NHP10*) (**Figure 2E-F**, **orange and pink**) and in the transcriptional regulator gene *TUP1* (**Figure 2E-F, orange**), which recruits the SAGA histone acetylase and SWI/SNF nucleosome-remodelling complex to stress-regulated promoters following Hog1 activation (Conaway & Conaway, 2009; Myer & Young, 1998; Proft & Struhl, 2002). Transposons in these genes likely disrupted the transcriptional reprogramming activities of Hog1 (Alepuz et al., 2003; Klopf et al., 2009; Mas et al., 2009; Nadal-Ribelles et al., 2012; Zapater et al., 2007), thereby suppressing the toxic effects of constitutive Hog1 activation. We did not observe any positive or negative selection for transposons inserted into genes previously proposed to act upstream of HHKs upon Fludioxonil treatment, such as genes affecting mitochondrial activity, glutathione homeostasis, or aldehyde metabolism (Brandhorst et al., 2019). This argues against the idea that Fludioxonil targets a factor upstream of group III HHKs. Together, this example demonstrates that SATAY can be performed in strains heterologously expressing antifungal drug targets, providing a straightforward way to screen antifungals with targets absent in *S. cerevisiae*.

### 3. Chitosan electrostatically interacts with cell wall mannosylphosphate

Having established that SATAY can identify antifungal resistance mechanisms and targets, we applied SATAY to examine Chitosan. Chitosan is a natural polysaccharide derived from the deacetylation of chitin (Kong et al., 2010) and is used to control numerous pre- and post-harvest fungal diseases (Bautista-Baños et al., 2006), but has an unclear mode of action. Chitosan’s activity is pH-dependent: at physiological pH, the amino groups in Chitosan are partially protonated (**Figure 3A**) (Ke et al., 2021) and probably interact electrostatically with the cell-surfaces of bacteria and fungi, increasing membrane permeability and causing metabolite leakage (Helander et al., 2001; Liu et al., 2004; Mellegård et al., 2011; Palma-Guerrero et al., 2010; Palmeira-de-Oliveira et al., 2011; Raafat et al., 2008; Tayel et al., 2010; Tsai & Su, 1999). However, the specific targets for Chitosan on fungal cell-surfaces have not been identified (Li et al., 2022; Ma et al., 2017; Shih et al., 2019).

**Figure 3.**
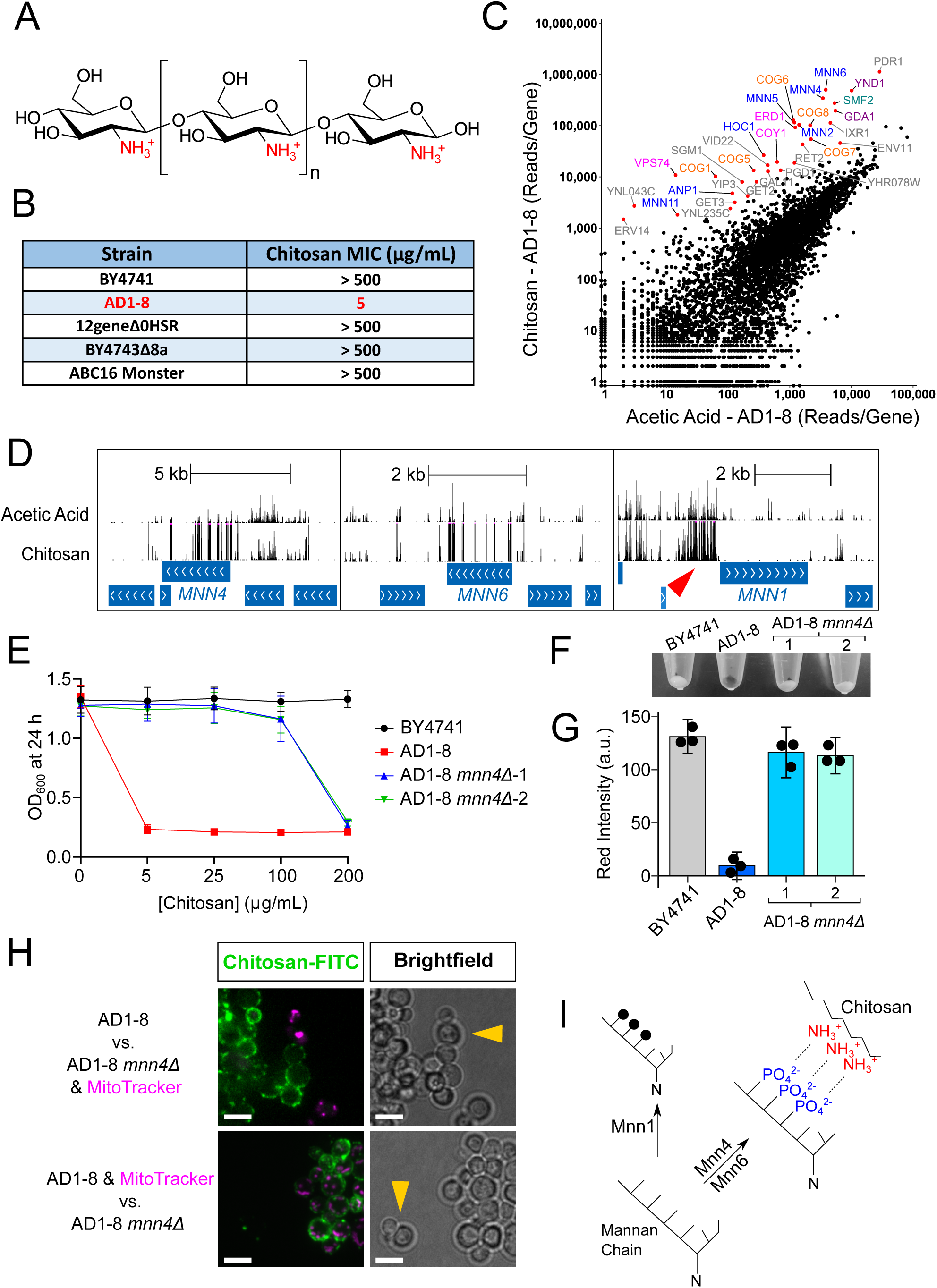
Chitosan electrostatically interacts with cell wall mannosylphosphate. **A)** Chemical structure of Chitosan. Protonated amino groups are shown in red. **B)** Minimal Inhibitory Concentrations (MICs) of Chitosan for five strains of *S. cerevisiae*. All strains are PDR-attenuated except for BY4741. MICs were determined after 24 hours for SC 2% glucose cultures incubated at 30°C, 200 rpm. Note in this figure, BY4741 refers to ByK352 (Supplementary Table 4). **C)** Scatter plot comparing the number of sequencing reads mapping to each of the 6603 yeast CDSs for an AD1-8 transposon library treated with acetic acid (0.0004%) or Chitosan (2 μg/mL). Gene names are colour-coded by biological process: blue – mannan chain elaboration on N-glycans of cell wall proteins; orange - CO G complex subunits; pink – intra-Golgi retrograde transport; purple – nucleoside phosphatases required for GDP-mannose transport into the Golgi; teal - manganese transport. **D)** Transposon insertion map for *MNN4, MNN6* and *MNN1* in the AD1-8 transposon library following treatments with acetic acid (0.0004% (v/v)) and Chitosan (2 μg/mL). The height of the bars corresponds to the number of sequencing reads, shown on a linear scale that is capped at 2,000. Horizontal blue bars represents CDSs. Red arrowhead indicates positively selected insertions in the promoter region of *MNN1*. **E)** Growth inhibition curves of a BY4741 strain, AD1-8, and two independent AD1-8 *mnn4*Δ strains treated with Chitosan in SC 2% glucose medium. OD_600_ measurements were recorded using a Bioscreen C™ instrument set at 30°C, continuous shaking. Each data point is the mean OD_600_ measurement for three technical replicates. Error bars show the standard deviation. **F)** Alcian blue staining of a BY4741 strain, AD1-8, and two independent AD1-8 *mnn4*Δ strains. Saturated cultures were treated with 10 μg/mL alcian blue solution in SC 2% glucose medium for 3 minutes. Image is displayed in the red channel, since alcian blue strongly absorbs red light. **G)** Quantification of background-subtracted red channel intensity of alcian blue-stained pellets of BY4741 strain, AD1-8, and two independent AD1-8 *mnn4*Δ strains. Three technical replicates for each strain. Error bars show the standard deviation. **H)** Representative images of AD1-8 and AD1-8 *mnn4*Δ strains following treatment with 25 μg/mL fluorescein-labelled Chitosan (green) for 2 hours in SC 2% glucose medium. Cells were treated in stationary phase. MitoTracker (magenta) was applied at 0.1 μM. Yellow arrowheads indicate representative AD1-8 *mnn4*Δ cells. Images are Z-stacks. Scale bars represent 5 μm. **I)** Model for the mechanism by which Chitosan binds to the yeast cell wall. Model depicts electrostatic interactions (black dashes) between Chitosan’s positively charged amino groups (red) and negatively charged mannosylphosphate groups (blue) on an N-linked glycan chain. Mannosylphosphate groups are added onto α1,2-linked mannose side branches by the mannosylphosphate transferases Mnn4 and Mnn6. Meanwhile, Mnn1 adds terminal α1,3-linked mannose residues (black circles) to α1,2-linked mannose side chains. Terminal α1,3-linked mannose residues compete with mannosylphosphate groups for available α1,2-linked mannose acceptor sites. *MNN1* overexpression in AD1-8 likely conferred Chitosan resistance by tipping this competition towards non-phosphorylated α1,3-linked mannose residues.

Despite having a broad antimicrobial spectrum (Goy et al., 2009; Ke et al., 2021), Chitosan has minimal activity against *S. cerevisiae* strains derived from the widely used laboratory strain S288C, such as BY4741 (Jaime et al., 2012; Zakrzewska et al., 2005, 2007). As yeast deletion collections were constructed in S288C (Giaever et al., 2002; Winzeler et al., 1999), chemogenomic studies using yeast deletion collections have evaluated Chitosan’s mode of action at high concentrations, limiting physiological and agricultural relevance (Galván Márquez et al., 2013; Jaime et al., 2012; Zakrzewska et al., 2007). However, as SATAY is not tied to a specific genetic background (Michel et al., 2017), we assessed the sensitivity of several *S. cerevisiae* strains to Chitosan. Chitosan susceptibility was observed for AD12345678 (hereafter AD1-8) (**Figure 3B**). AD1-8 is derived from a non-conventional strain called US50-18C, and lacks seven ABC transporters and the transcription factor *PDR3* (Supplementary Table 4) (Decottignies et al., 1998). Notably, S288C-derived strains lacking the same ABC transporters as AD1-8 (Chinen et al., 2011; Hoepfner et al., 2014; Suzuki et al., 2011) were not sensitive to Chitosan (**Figure 3B**), indicating that AD1-8’s Chitosan sensitivity was not attributable to the disruption of the PDR network, but to unknown characteristics of the US50-18C background.

We treated an AD1-8 transposon library with 2 μg/mL Chitosan and the corresponding acetic acid control. Chitosan caused a striking enrichment of transposons in genes associated with the GO terms “glycoprotein biosynthetic process” and “Golgi vesicle transport” (Supplementary Table 3). These included genes encoding Golgi-localised mannosyltransferases or mannosylphosphate transferases (**Figure 3C, blue**), which are responsible for elongating and modifying N-linked glycans on cell wall proteins (Munro, 2001). Additionally, Chitosan positively selected transposons in *GDA1* and *YND1* (**Figure 3C, purple**), which encode homologous Golgi-localised nucleoside phosphatases required for the transport of GDP-mannose into the Golgi lumen (Berninsone et al., 1994; Gao et al., 1999). Chitosan also enriched transposons in the CDS of *SMF2* (**Figure 3C, green**), which encodes a manganese (Mn^2+^) transporter necessary for the delivery of Mn^2+^ to the Golgi (Luk & Culotta, 2001). Consistently, Mn^2+^ is a critical co-factor for Ynd1 and Golgi-localised mannosyltransferases (Gao et al., 1999; Kuhn et al., 1991; Nakajima & Ballou, 1975). There was also a significant enrichment of transposons in *VPS74*, *ERD1*, *COY1* and five of the eight subunits of the <u>c</u>onserved <u>o</u>ligomeric <u>G</u>olgi (COG) complex (**Figure 3C, pink and orange**). These genes participate in intra-Golgi retrograde transport, which recycles glycosyltransferases from *trans*-Golgi to *cis/medial*-Golgi compartments, thereby maintaining their steady-state distribution. This is critical for ensuring the distinct steps of glycosylation occur in the correct sequential order (Bruinsma et al., 2004; Pokrovskaya et al., 2011; Sardana et al., 2021; Shestakova et al., 2006). Thus, our SATAY screen strongly indicates that Chitosan sensitivity in AD1-8 is engendered by the mannosylation of N-linked glycans on cell wall proteins, consistent with Chitosan interacting with the fungal cell wall (Li et al., 2022; Tayel et al., 2010).

Mannan chains on cell wall proteins are modified with phosphate groups, giving a net negative charge to the yeast cell wall (Karson & Ballou, 1978). We hypothesised that Chitosan might electrostatically interact with mannosylphosphate. This hypothesis is supported by multiple findings in our SATAY screen. First, transposons disrupting genes required for mannosylphosphate transfer, *MNN4* (Odani et al., 1996) and *MNN6* (Karson & Ballou, 1978; Wang et al., 1997), were strongly enriched by Chitosan (**Figure 3C-E**). Second, several genes acting upstream of *MNN4* and *MNN6* contained strongly selected transposons, namely *ANP1*, *MNN11*, *HOC1*, *MNN2* and *MNN5* (**Figure 3C**) (Munro, 2001; Orlean, 2012). These upstream genes are necessary for the addition of α1,2-linked mannose side branches, which are the sole substrates of Mnn4 and Mnn6 (Karson & Ballou, 1978). Third, Chitosan positively selected transposons in the promoter region of *MNN1* (**Figure 3D, red arrowhead**, Supplementary Figure S3A) and negatively selected transposons in the CDS of this gene. Mnn1 adds terminal α1,3-linked mannose residues to α1,2-linked mannose side chains (Raschke et al., 1973; Yip et al., 1994). As such, terminal α1,3-linked mannose residues compete with mannosylphosphate groups for available acceptor sites (Karson & Ballou, 1978). Thus, *MNN1* overexpression likely conferred Chitosan resistance in AD1-8 by tipping this competition towards non-phosphorylated α1,3-linked mannose residues (**Figure 3I**).

Alcian blue is a cationic dye that can distinguish yeasts with different cell wall mannosylphosphate compositions (Ballou, 1990). AD1-8 cells were strongly stained by alcian blue, and this staining was dependent on *MNN4* (**Figure 3F-G**). In contrast, despite being wildtype for *MNN4*, the Chitosan-resistant lab strain BY4741 did not bind alcian blue. This shows that levels of cell wall mannosylphosphate correlated with Chitosan sensitivity, supporting the model that they interact electrostatically. Finally, our model implies that Chitosan sensitivity and resistance are determined by different abilities to bind Chitosan. To test this, we treated cells with fluorescein-labelled Chitosan. To directly compare the fluorescence staining intensities on the same images, strains were mixed pairwise after staining and then imaged. The strains were distinguishable because only one strain received a pre-treatment with the unrelated vital stain, MitoTracker. Fluorescein-labelled Chitosan strongly stained the cell walls of AD1-8 WT cells but failed to bind the cell walls of AD1-8 *mnn4*Δ cells (**Figure 3H**) or BY4741 cells (Supplementary Figure S7). This further indicates that Chitosan exerts its fungistatic effect by binding mannosylphosphate on the cell-surface (**Figure 3I**). Our study therefore identifies a mode of action for Chitosan on fungal cell walls.

### 4. Hol1 imports ATI-2307, Pentamidine and Iminoctadine into *S. cerevisiae*

ATI-2307 (formerly T-2307) is a novel antifungal with an unclear mode of action (Wiederhold, 2021). ATI-2307 exhibits broad-spectrum *in vitro* and *in vivo* activity against many clinically important fungal pathogens, including Fluconazole-resistant strains of *C. albicans* and echinocandin-resistant *Candida spp.* (Mitsuyama et al., 2008; Wiederhold et al., 2020; Nishikawa et al., 2017; Wiederhold et al., 2015, 2016). ATI-2307 therefore represents a promising compound for the future control of fungal infections.

ATI-2307 specifically inhibits *S. cerevisiae* in non-fermentative growth conditions (Shibata et al., 2012). Upon challenging a wild-type transposon library with ATI-2307 in non-fermentative medium, we identified multiple genes affecting susceptibility to this compound. Notably, genes causing ATI-2307 hypersensitivity when interrupted by transposons were significantly associated with the GO term “Regulation of Gluconeogenesis” (Supplementary Table 3). This included subunits of the GID (<u>G</u>lucose Induced degradation <u>D</u>eficient) complex, namely *GID7, GID8, VID28* and *VID30* (Supplementary Figure 1A, top-right panel). GID is an E3 ligase that polyubiquitylates fructose-1,6-bisphosphatase when non-fermenting cells encounter glucose (Regelmann et al., 2003). *UBC8* was also identified, which encodes a ubiquitin-conjugating enzyme that regulates the activity of the GID complex (Regelmann et al., 2003; Schule et al., 2000). Moreover, transcription factors regulating the expression of gluconeogenic genes were essential on ATI-2307, namely *RDS2* (Soontorngun et al., 2007) and *ERT1* (Gasmi et al., 2014) (Supplementary Figure 1A, top-right panel). The connection between gluconeogenesis and ATI-2307 susceptibility is unclear, but the enrichment of these functionally-related hits underlines the specificity and comprehensiveness of the screen.

Among the genes causing ATI-2307 resistance when disrupted by transposons were the paralogs *RMD1* and *RMD8* (**Figure 4A, green**). Very little is known about the function of these genes, but they are homologous to a third protein, Mrx10, an inner mitochondrial membrane component of MIOREX complexes, large assemblies that spatially organise mitochondrial protein synthesis (Kehrein et al., 2015). Although Mrx10 was not identified by the SATAY screen, the homology to Rmd1 and Rmd8 is notable because mitochondrial protein synthesis is required for the biogenesis of respiratory complexes, which were previously implicated as targets for ATI-2307 (Yamashita et al., 2019).

**Figure 4.**
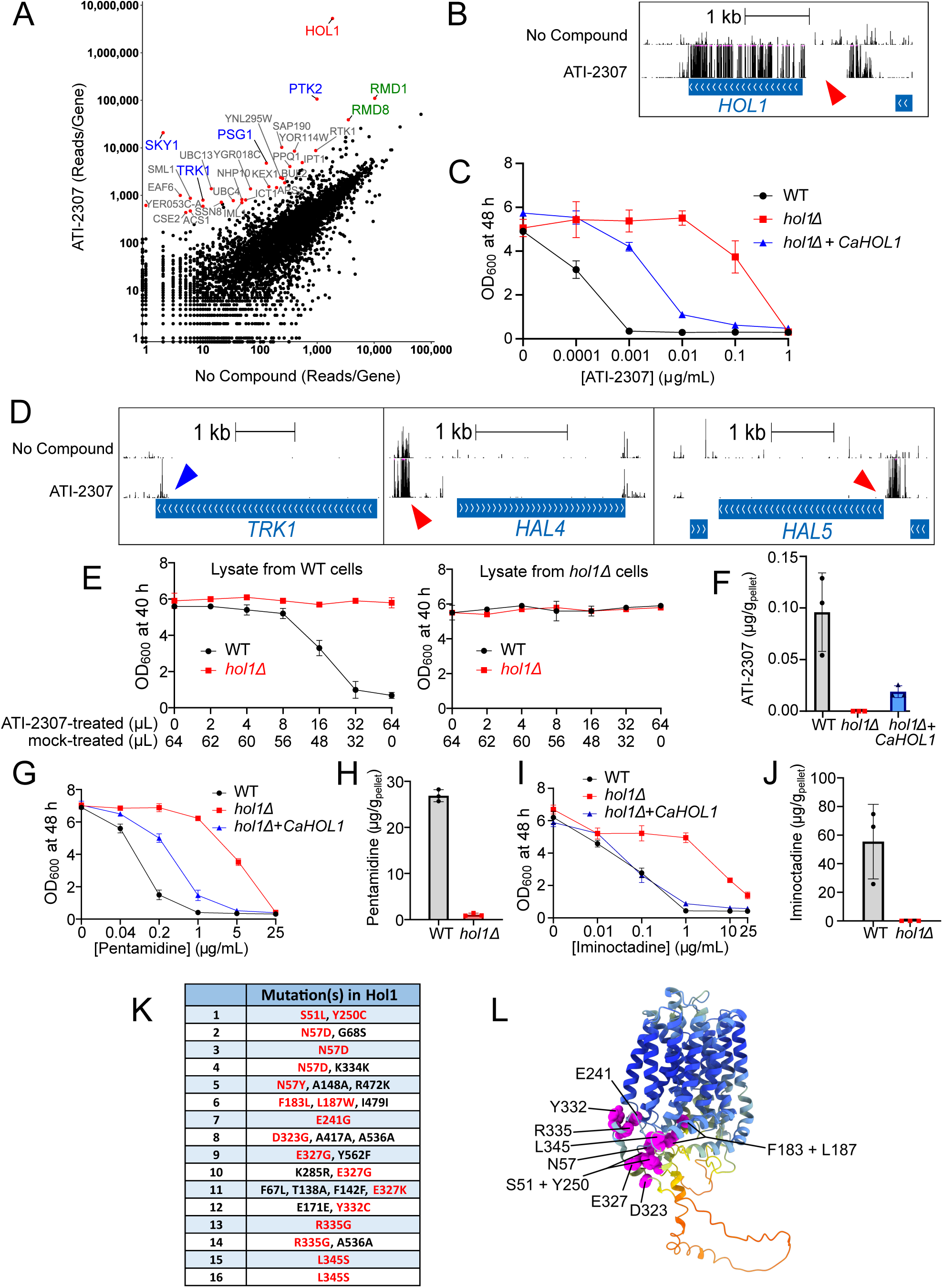
Hol1 imports ATI-2307, Pentamidine and Iminoctadine into *S. cerevisiae*. **A)** Scatter plot comparing the number of sequencing reads mapping to each of the 6603 yeast CDSs in a transposon library after a control treatment versus ATI-2307 treatment (0.000175 μg/mL) in SC -TRP 2% ethanol medium. *HOL1* is coloured red. Genes involved in the regulation of the proton-motive force are coloured blue. *RMD1* and *RMD8* are coloured green. **B)** Transposon insertion map for *HOL1* for the control and ATI-2307 treatments. The height of the bars corresponds to the number of sequencing reads, shown on a linear scale that is capped at 500. Red arrowhead indicates region upstream of *HOL1* where transposon insertions were negatively selected by ATI-2307. This region negatively regulates *HOL1* expression in a post-transcriptional manner (Vindu et al., 2021). **C)** Growth inhibition curves for wild-type, *hol1*Δ and *hol1*Δ *CaHOL1* strains treated with ATI-2307. WT and *hol1*Δ strains were transformed with pRS316 empty vector. OD_600_ measurements were recorded after 48 hours of growth in SC -URA 2% ethanol medium at 30°C, 200rpm. Each data point is the mean OD_600_ measurement for three independent biological replicates. Error bars show the standard deviation. Error bars for some measurements were too small to be displayed. **D)** Transposon insertion maps for *TRK1*, *HAL4* and *HAL5* after the control and ATI-2307 treatments. The height of the bars corresponds to the number of sequencing reads, shown on a linear scale that is capped at 500. Blue arrowhead indicates positively selected insertions in the 3’-end of *TRK1*. Red arrowheads indicate positively selected insertions in the promoter regions of *HAL4* and *HAL5*. **E)** Growth inhibition curves for WT and *hol1*Δ strains treated with lysate derived from ATI-2307-treated-WT or *hol1*Δ cells. OD_600_ measurements were recorded after 40 hours of growth in SC 2% ethanol medium at 30°C, 200rpm. Each data point is the mean of three independent biological replicates. Each culture was inoculated with a total of 64 μL lysate derived from WT or *hol1*Δ cells. The volume of lysate from ATI-2307-treated versus mock-treated cells is indicated on the x-axis. Error bars show the standard deviation. **F)** LC-MS analysis of ATI-2307 uptake in WT, *hol1*Δ and *hol1*Δ *CaHOL1* cells. Cells were cultured in 200 mL SC 2% ethanol at 30°C, 200rpm from a starting OD_600_ 0.5. Upon reaching OD_600_ 1, cells were treated for 2 hrs with ATI-2307 (0.003 μg/mL). The *hol1*Δ strains expressing *CaHOL1* (pBK994) were grown SC -URA 2% ethanol medium. Quantifications of ATI-2307 were performed using matrix-matched calibration curves. Each dot represents an independent biological replicate (n = 3). Error bars show the standard deviation. **G)** Growth inhibition curves for wild-type, *hol1*Δ and *hol1*Δ *CaHOL1* strains treated with Pentamidine, as in **C)**. **H)** LC-MS analysis of Pentamidine uptake in WT or *hol1*Δ cells. Cells were treated with 1 μg/mL Pentamidine. Other experiment details are identical to those described for ATI-2307 in **F)**. **I)** Growth inhibition curves for wild-type, *hol1*Δ and *hol1*Δ *CaHOL1* strains treated with Iminoctadine, as in **C)**. **J)** LC-MS analysis of Iminoctadine uptake in WT or *hol1*Δ pellets. Cells were treated with 1 μg/mL Iminoctadine. Other experiment details are identical to those described for ATI-2307 in **F)**. **K)** Mutations identified in Hol1 from sixteen *spe1*Δ *hol1*Δ p[*HOL1*] strains exhibiting a separation-of-function phenotype. For each strain, all the mutations identified in Hol1 are listed, including synonymous mutations. Non-synonymous mutations shown in red were considered causal of the separation-of-function phenotype. For plasmids bearing multiple non-synonymous mutations in Hol1, the causal mutation was assigned to mutations identified in several independent plasmids e.g. N57 and E327. **L)** AlphaFold2 model of the *S. cerevisiae* Hol1 protein (UniProt Identifier P53389) showing the positions of the amino acid residues that were considered causal of the separation-of-function phenotype when mutated.

The CDS of *HOL1* contained the most strongly selected transposon insertions (**Figure 4A-B**). Additionally, transposons mapping upstream of *HOL1*, in a region that negatively regulates *HOL1* expression in a post-transcriptional manner (Vindu et al., 2021; Wright et al., 1996), were negatively selected by ATI-2307 (**Figure 4B, red arrowhead**). These findings indicate that Hol1 engenders ATI-2307 susceptibility. Hol1 was recently identified as a high-affinity polyamine transporter in *S. cerevisiae* (Vindu et al., 2021). Strikingly, ATI-2307 uptake by *C. albicans* depends on the activity of an unidentified polyamine transporter (Nishikawa et al., 2010, 2016). Together, this suggested that Hol1 might be the transporter responsible for ATI-2307 uptake. Consistently, the deletion of *HOL1* confers ATI-2307 resistance, increasing the minimal inhibitory concentration (MIC) 1,000-fold from 0.001 to 1 µg/mL (**Figure 4C**). Moreover, Hol1 transport is driven by the plasma membrane proton-motive force (PMF) (Vindu et al., 2021), and transposon insertions predicted to weaken the PMF were positively selected by ATI-2307. This included insertions in the CDSs of *PTK2* and *PSG1* (**Figure 4A, blue**), both of which positively regulate the H^+^-ATPase Pma1 (Eraso et al., 2006; Geva et al., 2017; Goossens et al., 2000) which generates the PMF (Serrano et al., 1986). ATI-2307 also positively selected transposons mapping to the extreme 3’-end of *TRK1* (**Figure 4D, blue arrowhead**). The deletion of the last 35 amino acids of Trk1 increases the stability of the Trk1-Trk2 potassium ion transporter (Pérez-Valle et al., 2007) which dissipates the PMF (Gaber et al., 1988; Ko et al., 1990; Madrid et al., 1998). There was also a significant enrichment of transposons in the promoters of *HAL4* and *HAL5* (**Figure 4D, red arrowheads**), two partially redundant positive regulators of Trk1-Trk2, and in the CDS of *SKY1* (**Figure 4A, blue**), a negative regulator of Trk1-Trk2 (Mulet et al., 1999; Pérez-Valle et al., 2007; Forment et al., 2002). Like *ptk2*Δ mutants, *sky1*Δ mutants are resistant to toxic concentrations of ATI-2307 (Nishikawa et al., 2016) and polyamines (Erez & Kahana, 2001; Forment et al., 2002; Kaouass et al., 1997).

ATI-2307 accumulates to high concentrations inside *C. albicans* cells (Nishikawa et al., 2010). We reasoned that if the same was true in *S. cerevisiae*, lysate from ATI-2307-treated WT cells may contain enough ATI-2307 to inhibit the growth of another *S. cerevisiae* culture, and that this might depend on Hol1. We treated non-fermentative WT and *hol1*Δ cultures in log-phase with an inhibitory concentration of ATI-2307 (0.003 μg/mL) for two hours. Cells were then harvested, washed, and homogenised. Lysate from ATI-2307-treated WT cells inhibited the growth of non-fermentative WT cultures, but not *hol1*Δ cultures (**Figure 4E, left**), indicating that the growth inhibition was indeed due to ATI-2307. In contrast, lysate from ATI-2307-treated *hol1*Δ cells lacked inhibitory activity (**Figure 4E, right**), consistent with Hol1-dependent ATI-2307 accumulation in *S. cerevisiae*. Liquid Chromatography – Mass Spectrometry (LC-MS) was then used to quantify Hol1-dependent ATI-2307 accumulation in WT and *hol1*Δ cells. Whilst an average of 0.1 μg ATI-2307 was detected per gram of WT pellet, the concentration in *hol1*Δ lysates was below the detection limit (i.e. less than ∼0.005 μg/mL) (**Figure 4F**). Thus, Hol1 is responsible for the selective accumulation of ATI-2307 in *S. cerevisiae*. Moreover, the heterologous expression of *C. albicans HOL1* (*CaHOL1*) partially restored ATI-2307 uptake (**Figure 4F**) and sensitivity (**Figure 4C**) in the *hol1*Δ mutants, indicating that Hol1 also imports ATI-2307 into fungal pathogens.

Pentamidine is an antiprotozoal drug that shares a similar chemical structure to ATI-2307 (Supplementary Figure S8). Given that Pentamidine competitively inhibits the uptake of polyamines and ATI-2307 by *C. albicans* (Nishikawa et al., 2010, 2016), we hypothesised that Hol1 also imports Pentamidine. Consistent with this, *hol1*Δ mutants showed reduced Pentamidine susceptibility (**Figure 4G**) and uptake (**Figure 4H**), and Pentamidine susceptibility was partially restored by the heterologous expression of *CaHOL1* (**Figure 4G**). Additionally, we assessed whether Hol1 imports Iminoctadine (Supplementary Figure S8), a polyamine found within the fungicide Guazatine (Castagnolo et al., 2011) that has activity against multiple fungal phytopathogens (Ryu et al., 2014; Zhu et al., 2019) and fluconazole-resistant *Candida spp*. (Castagnolo et al., 2011). As shown in **Figure 4I**, Iminoctadine susceptibility was reduced by the deletion of *HOL1*, which could be reversed by *CaHOL1* expression. Consistently, WT and *hol1*Δ cells treated for two hours with 1µg/mL Iminoctadine showed a striking 3,500-fold difference in Iminoctadine uptake (52.5µg per gram of WT pellet versus 0.015µg per gram of *hol1*Δ pellet) (**Figure 4J**). These results demonstrate that Hol1 also imports Pentamidine and Iminoctadine.

Our results indicate that loss-of-function mutations in Hol1 could confer ATI-2307 resistance in the clinic. However, the complete loss of this polyamine transporter is likely to carry a high fitness cost in low-nitrogen conditions (Vindu et al., 2021). This fitness cost might be circumvented by point mutations in *HOL1* that prevent ATI-2307 uptake while preserving polyamine import. Such separation-of-function mutations could present a caveat for the continued therapeutic development of ATI-2307. To assess whether separation-of-function mutations can arise, we performed a forward genetics screen that coupled random PCR mutagenesis to phenotypic screening. The principle underpinning the screen is that *HOL1* is essential for the growth of the polyamine auxotrophic *spe1*Δ strain in low-polyamine conditions (e.g. SC 2% ethanol + 1 μM spermidine, Supplementary Figure S12), but not in rich medium (Fonzi, 1989; Whitney & Morris, 1978). A plasmid library of randomly mutagenized *HOL1* genes was generated by gap-repair cloning in a *spe1*Δ *hol1*Δ strain. Approximately 10,000 transformants were then replica plated onto SC 2% ethanol medium containing 0.01 μg/mL ATI-2307 and 1 μM spermidine. Sixteen transformants showing ATI-2307 resistance were isolated before the plasmids were rescued. Seven plasmids were transformed back into the *spe1*Δ *hol1*Δ strain and disc diffusion assays confirmed resistance to ATI-2307, Pentamidine and Iminoctadine (Supplementary Figure S13). Given that *HOL1* is essential for *spe1*Δ strains on SC 2% ethanol (Supplementary Figure S12), the resistance phenotypes confirmed that the *HOL1* alleles allowed spermidine uptake but limited import of ATI-2307, Pentamidine and Iminoctadine. These results indicate that the transport activities of Hol1 can be functionally separated. All sixteen plasmids were sequenced and contained at least one non-synonymous mutation in *HOL1* (**Figure 4K**). Furthermore, the codons for N57, L325, and E327 were mutated in more than one independent plasmid, suggesting that our screen was approaching saturation. Mapping the mutated residues onto the AlphaFold2 model of Hol1 (Jumper et al., 2021) showed that most of these mutations clustered on the cytoplasmic side of Hol1 near the base of the pore (**Figure 4L**). This clustering suggests that our screen did not merely identify partial loss-of-function mutations in *HOL1*, but instead pinpointed a specific domain critical in mediating ATI-2307 uptake. Therefore, Hol1 mutations can disrupt its antifungal transport function without sacrificing its ability to transport polyamines, potentially allowing antifungal resistance to develop with minimal fitness cost.

## Discussion

### 1. SATAY can accelerate the discovery of antifungal resistance mechanisms and modes of action

The rapid emergence of multidrug-resistant fungal pathogens is undermining our ability to control fungal infections. This problem is exacerbated by our incomplete understanding of the molecular mechanisms governing antifungal resistance and our limited armamentarium of antifungal compounds. Here, SATAY is applied to reveal loss- and gain-of-function mutations conferring resistance and sensitivity to 20 different antifungal compounds. These mutations pinpoint antifungal drug targets, transporters responsible for drug uptake or drug efflux, as well as metabolic, signalling and trafficking pathways affecting antifungal susceptibility. This study therefore provides a comprehensive dataset of cellular mechanisms that buffer fungal cells against diverse chemical stresses, which could inform the design of combination therapies and resistance management strategies. Moreover, by performing SATAY in PDR-attenuated strains, strains heterologously expressing drug targets, and non-conventional strains, we identify resistance mechanisms and targets for compounds that lack activity against conventional laboratory strains of *S. cerevisiae*. SATAY therefore augments the range of antifungals amenable for screening with *S. cerevisiae*, creating opportunities for determining how resistance emerges against a broader spectrum of compounds.

### 2. Chitosan electrostatically interacts with mannosylphosphate on fungal cell-surfaces

Chitosan is a natural antifungal compound with several attractive qualities, including biocompatibility, biodegradability, and non-toxicity. However, its application is constrained by the fact acidic conditions are required for solubility. Efforts are underway to create Chitosan derivatives that improve bioavailability while preserving antifungal efficacy (Hu & Luo, 2016; Verlee et al., 2017). One key challenge in these efforts is our lack of detailed knowledge regarding Chitosan’s specific molecular targets in fungi (Qin & Li, 2020). This gap has persisted partly because widely employed laboratory strains of *S. cerevisiae* such as S288C, which are used in conventional chemogenomic screening approaches, have minimal sensitivity to Chitosan. Here, we perform SATAY in the Chitosan-sensitive strain AD1-8, derived from the non-conventional strain US50-18C (Decottignies et al., 1998), and uncover mannosylphosphate as the cell wall target for Chitosan. This finding may guide the development of Chitosan derivatives with improved solubility and antifungal activity.

Interestingly, S288C-derived strains lacked cell wall mannosylphosphate, even though they harbour wild-type copies of *MNN4* and *MNN6*. The genetic determinants underpinning Chitosan resistance in S288C-derived strains are multi-factorial, since upon crossing AD1-8 to ByK352 (an S288C-derived strain), we observed that Chitosan sensitivity and alcian blue staining were inherited in a polygenic fashion (data not shown). The cell-surfaces of S288C-derived strains have been modified by decades of selection in laboratory environments. For instance, S288C strains were deliberately selected to be defective in flocculation (Mortimer & Johnston, 1986), a process leading to unwanted cellular aggregations (Stratford, 1992). This is notable because mannan chains on the yeast cell wall are required for flocculation (Stratford, 1992). It seems possible that yeast geneticists have inadvertently selected for other mutations that affected cell wall mannosylphosphate. Additional basic traits for fungal pathogenicity, such as pseudo-hyphal growth (Gancedo, 2001) and biofilm formation (Bojsen et al., 2012), might have been accidentally or purposely lost from *S. cerevisiae* laboratory strains. SATAY therefore offers a unique opportunity for screening compounds targeting such processes in non-deficient, wild strains.

### 3. Hol1 imports ATI-2307, Pentamidine and Iminoctadine into *S. cerevisiae*

We have identified that Hol1 is the high-affinity polyamine transporter responsible for the uptake and accumulation of ATI-2307, Pentamidine and Iminoctadine in *S. cerevisiae*. Previous studies have reported that ATI-2307 has broad-spectrum activity against clinically significant *Candida* species, including *C. albicans*, *C. glabrata*, *C. parapsilosis*, *C. tropicalis* and *C. auris* (Mitsuyama et al., 2008; Wiederhold, et al., 2020). Given that Hol1 from *C. albicans* can import ATI-2307 and that Hol1 is highly conserved across *Candida* species (Supplementary Figure S14), it seems plausible that Hol1 orthologs across the *Candida* clade import ATI-2307.

Although carrier-mediated import of antifungal compounds has been described previously (Galocha et al., 2020; Lanthaler et al., 2011), the propensity of Hol1 to concentrate ATI-2307 is unique and underpins the potency of this novel antifungal against yeasts. Few studies have investigated drug uptake by fungal pathogens (Galocha et al., 2020), yet they might harbour broad-spectrum, high-affinity nutrient permeases that could be hijacked by novel antifungal compounds. Identifying and characterising such transporters could be a promising strategy for bolstering antifungal development pipelines. The promiscuity of Hol1, with only loose structural homology between its identified substrates (Supplementary Figure 8), makes it a promising candidate for the development of actively imported drugs or pro-drugs.

However, our study also reveals a straightforward pathway for the emergence of ATI-2307 resistance. While Hol1 is essential for polyamine uptake (Vindu et al., 2021), the single point mutations that we identified, which limit ATI-2307 uptake without perturbing polyamine uptake, could confer ATI-2307 resistance with little fitness cost. Gradual accumulation of additional mutations through sustained selection in the clinic might reduce any remaining costs. Therefore, by demonstrating that there exists a straightforward pathway for ATI-2307 resistance to arise, our findings may support further precautionary measures in the clinical use of ATI-2307.

## Materials and Methods

### Yeast strains and culture conditions

*S. cerevisiae* strains and primers are listed in Supplementary Tables 4 and 5, respectively. Gene deletions were generated by PCR-mediated gene disruption with selectable markers using the Longtine and Janke toolboxes (Janke et al., 2004; Longtine et al., 1998). Yeast transformations were performed using lithium acetate (Gietz & Schiestl, 2007). Yeast strains yBL152 (ByK831, W303-1B), 12geneΔ0HSR (ByK1151), AD12345678 (ByK1174), BY4743Δ8a (ByK1230) and ABC16 Monster (ByK1266) were received as kind gifts from Prof. Mattias Peter (ETH Zurich), Prof. Françoise Strutz (University of Geneva), Prof. Mohan Gupta (University of Iowa), Prof. Martin Spiess (University of Basel), Prof. Frederick Roth (University of Toronto), respectively.

Media recipes are described in the Supplementary Methods. Fermentative cultures were grown in media prepared using amino acid recipe A, and non-fermentative cultures were grown in media prepared using amino acid recipe B. All cultures were incubated at 30°C, 200 rpm.

### Plasmids

Plasmids are listed in Supplementary Table 6. Restriction enzymes were purchased from NEB.

To construct the plasmid p405-*TEF1*pr-*BdDRK1* (pBK864), the *DRK1* CDS from *Blastomyces dermatitidis* (NCBI Reference Sequence: EGE84246.1) was commercially synthesised with yeast codon optimisation, 5’ restriction site for *Spe*I and 3’ restriction site for *Pst*I (Biomatik, Canada). The *BdDRK1* CDS was then sub-cloned into p405-*TEF1*pr (Addgene plasmid #15968) using *Spe*I and *Pst*I and T4 DNA ligase (Thermo Fisher Scientific #EL0011).

The plasmid bearing the *HOL1* CDS from *Candida albicans* (pBK994) was generated by gap-repair cloning. Briefly, the *CaHOL1* CDS was PCR-amplified from genomic DNA derived from *Candida albicans* strain ATCC MYA-2876 using primers 11 and 12 (Supplementary Table 5). The 5’ and 3’ ends of the *CaHOL1* PCR product were homologous to the *ScHOL1* 5’ UTR and 3’ UTR sequences on pBK958. The *ScHOL1* CDS was excised from pBK958 using *Bsu36*I and *Nru*I. The linearised pBK958 backbone was co-transformed with the *CaHOL1* PCR product into *hol1*Δ strain of *S. cerevisiae* (ByK2032). Plasmid pBK994 was verified by colony PCR.

### Antifungal compounds

Antifungal compounds are listed in Supplementary Table 2. Compounds were stored at the manufacturer’s recommended temperature. Stocks were prepared using dimethyl sulfoxide (DMSO) (Fisher Scientific, 10103483) or Milli-Q^®^ and stored at - 20°C for maximally 18 months.

### Chitosan preparation

Stocks of medium molecular weight Chitosan (190-310 kDa, 75-85% deacetylated, shrimp shells) (Sigma-Aldrich, 448877) were prepared by nitrous deamination, as described previously (Zakrzewska et al., 2005). Briefly, 10 g Chitosan was dissolved in 200 mL 10% acetic acid (Sigma-Aldrich, A6283) and incubated with 200 mg sodium nitrite (Sigma-Aldrich, 237213) in the dark at room temperature for 17 hrs. The reaction was terminated by adjusting to pH 5.5 with 5M NaOH.

### Antifungal susceptibility tests

#### Bioscreen C

Sub-lethal concentrations of antifungal compounds were determined using a Bioscreen C^™^ instrument (Oy Growth Curves Ab Ltd, Helsinki, Finland). Cells were cultured from OD_600_ 0.2 in SC 2% glucose medium containing antifungal compounds at 30°C for 24 hrs with continuous shaking, and OD_600_ readings were recorded every 20 minutes. Final DMSO concentrations in the cultures never exceeded 1% (v/v).

#### ATI-2307, Pentamidine and Iminoctadine susceptibility tests

Yeast strains grown to saturation in SC -URA 2% ethanol 0.2% glucose were reseeded at OD_600_ 0.2 in SC -URA 2% ethanol containing ATI-2307, Pentamidine or Iminoctadine. Wild-type strains were transformed with an empty CEN/*URA3* plasmid (pRS316) (Sikorski & Hieter, 1983) whilst *hol1*Δ strains were transformed with pRS316 or plasmid pBK994, encoding *HOL1* from *Candida albicans*.

### SAturated Transposon Analysis in Yeast (SATAY)

#### Library generation on SC -ADE 2% galactose plates

Transposon libraries were generated on SC -ADE 2% galactose plates using previously described protocols (Michel et al., 2017; Michel & Kornmann 2022), with the following modifications. First, the OD_600_ at plating for strains 12geneΔ0HSR *ade2*Δ (ByK1261) and AD12345678 *ade2*Δ (ByK1262) was 65 and 160, respectively. Second, the AD12345678 *ade2*Δ transposon library was harvested ten days after transposition induction. Third, transposon libraries were not expanded in SC -ADE 2% glucose after harvest.

#### Library generation in liquid media

The culture volumes and media used for the generation of transposon libraries in liquid medium are summarised in Supplementary Table 7. Yeast strains were transformed with either pBK549 or pBK626. Individual pBK549 transformants were streaked on SC -URA 2% glucose and SC -ADE 2% glucose plates. For transformants that showed full growth on SC - URA whilst producing only a few colonies (2-6) on SC -ADE were scraped from the SC -URA plates and used to inoculate pre-cultures of SC -URA 2% raffinose 0.2% glucose at OD_600_ 0.2 and grown to saturation. To induce transposition, saturated pre-cultures were diluted to OD_600_ 0.2 in SC -URA 2% galactose and grown for ∼50 hours at 30°C, 200 rpm. Transposed cells were then diluted in SC -ADE 2% glucose and grown for ∼70h to OD_600_ ∼2. Dilutions of transposed cultures were plated at T_0_, T_20_ and at the end of induction on SC, SC -URA, and SC -ADE plates. Plating at T_0_ served to estimate the background of ADE+ clones due to spontaneous recombination of pBK549 (typically 0.01% in BY4741 background and 0.007% in W303 background). Plating at T_20_ guided the re-inoculation following transposition. The number of ADE+ cells at T_20_ typically corresponds to ∼10% of the number of ADE+ cells at the end of induction.

The protocol for generating transposon libraries with pBK626 transformants included the following modifications. First, individual pBK626-transformants were streaked on SC -URA 2% glucose and SC -TRP 2% glucose. Second, dilutions of induced cultures were plated at T_0_, T_20_ and ∼T_50_ on SC, SC -URA, and SC -TRP plates. Third, following transposition, cells were diluted in SC -TRP 2% glucose at OD_600_ 1 and grown to saturation, before a second re-growth was performed in SC -TRP 2% glucose from OD_600_ 0.3 to saturation. Fourth, cells were pelleted and stored at -80°C as 30% (v/v) glycerol stocks.

Prior to ATI-2307 treatment, the ByK831 transposon library was thawed from -80°C glycerol stock and grown from OD_600_ 0.25 to saturation in 1.2L of SC -TRP 2% glucose.

#### Antifungal compound treatments of transposon libraries

Transposon libraries were treated for two successive growth rounds from OD_600_ 0.1-0.2 to saturation (OD_600_ ∼5) at 30°C, 200rpm in the presence of antifungal compounds at the concentrations indicated in Supplementary Table 2, in media and volumes summarised in Supplementary Table 8. Cultures were harvested for cell pellets after the second growth round, using previously described protocols (Michel et al., 2017; Michel & Kornmann 2022) and stored at -80°C.

The first treatments with Amphotericin B (0.75 μg/mL) and Caspofungin (0.03 μg/mL) initially prevented growth. These cultures were diluted 2X in fresh medium after 24 hrs. The first treatment of Oxytriazine (1.491 μg/mL) treatment prevented growth so culture was diluted 3X in fresh medium after 46 hrs.

Prior to ATI-2307 treatment in SC -TRP 2% ethanol, the ByK831 transposon library was adapted to respiratory conditions by growing to saturation in 500 mL SC -TRP 2% ethanol 0.2% glucose medium.

#### Transposon library sequencing and bioinformatics analysis

Cell pellets were processed for sequencing following previously described protocols (Michel et al., 2017; Michel & Kornmann 2022). Barcoded transposon libraries were pooled and sequenced using the NextSeq^TM^ 500/550 High Output Kit v2.5 (75 cycles) (Illumina, 20024906) and NextSeq^TM^ 500 Sequencing System (Illumina, SY-415-1001) following the manufacturer’s guidelines. Bioinformatics analysis of SATAY sequencing data was performed as described in (Michel et al., 2017; Michel & Kornmann 2022). The *.bed* and *.wig* output files were uploaded to UCSC Genome Browser to visualise transposition events and read counts per transposon, respectively.

### Gene Ontology term analyses

Gene Ontology (GO) term analyses were performed using GO Term Finder (v0.86) (https://www.yeastgenome.org/) for the 25 – 50 genes with the highest read counts (or the lowest number of transposons) in the compound treatment relative to the control treatment. The background set of genes consisted of all the 6603 genes in the *S. cerevisiae* genome. Holm-Bonferroni-corrected GO term enrichments were computed for the “Process” Ontology aspect with a *p*-value below 0.01.

### Alcian blue dye binding assay

Saturated yeast grown in SC 2% glucose were treated with 10 μg/mL Alcian blue solution (Sigma-Aldrich, 1016470500) at room temperature for 3 minutes after brief vortexing. Cells were then centrifuged before the pellets were imaged.

### Live imaging of yeast

Yeast grown to saturation in SC 2% glucose medium were treated with 25 μg/mL fluorescein-labelled Chitosan (CD Bioparticles, CDHA061) for 2 hours. For the last 30 minutes of the fluorescein-labelled Chitosan treatment, cells were optionally treated with 0.1 μM MitoTracker™ Orange CMTMRos (Thermo Fisher Scientific, M7510). Cells were then washed twice and mixed pairwise, transferred to the microscope slide and imaged using an IX81 Olympus® inverted spinning disk microscope with an EM-CCD camera (Hamamatsu Photonics) using a 100× oil objective (NA = 1.4). Images were analysed in ImageJ (v1.53c).

### Assessing ATI-2307, Pentamidine and Iminoctadine accumulation in *S. cerevisiae*

#### Generating cell lysate

Wild-type and *hol1*Δ strains were grown to saturation in SC 2% ethanol 0.2% glucose. Cells were then reseeded at OD_600_ 0.5 in SC 2% ethanol (1L for subsequent cell lysate treatments; 200 mL for subsequent LC-MS experiments), and upon reaching OD_600_ ∼1, the cells were treated with ATI-2307 (0.003 μg/mL), Pentamidine (1 μg/mL) or Iminoctadine (1 μg/mL) for 2 hrs. Cultures were harvested by centrifugation before cell pellets were washed three times in SC 2% ethanol.

For subsequent yeast growth inhibition, cell lysate was obtained by resuspending ∼500 mg cell pellets in 500 μL Cell Breaking Buffer (2% Triton X-100, 1% SDS, 100 mM NaCl, 100 mM Tris–HCl, pH 8.0, 1 mM EDTA). Cell suspensions were distributed into 280 μL aliquots, to which 400 μL SC 2% ethanol and 200 μL 0.5 mm Glass Mill Beads (BioSpec Products, 11079105) were added, before vortexing at 2850 rpm for 10 minutes, 4°C, using a Disruptor Genie (Scientific Industries, SI-DD38). Aliquots were centrifuged before the lysate was collected.

For LC-MS, ∼500 mg cell pellets were resuspended in 500 μL 80% (v/v) acetonitrile (HPLC-grade, VWR) 20% (v/v) Milli-Q^®^ and stored at -20°C for ∼24 hours. Cell suspensions were then transferred to 2 mL tubes (Precellys^®^ MK28-R Hard Tissue Grinding Kit, Bertin Technologies), from which the steel beads were replaced with 200 μL 0.5 mm Glass Mill Beads (BioSpec Products, 11079105). Bead-beating was performed using a Precellys® Evolution homogeniser (Bertin Technologies, P002511-PEVT0-A.0) at 6,000 rpm for 10 cycles of: 1-minute beating, 30 second pause, 1-minute beating. Cycles were spaced by 2-minute intervals, during which the samples were kept on ice. Samples centrifuged before the lysates were collected and stored at -20°C until LC-MS analysis.

#### Cell lysate treatments

Yeast grown to saturation in SC 2% ethanol 0.2% glucose were reseeded at OD_600_ 0.2 in 10mL SC 2% ethanol and treated with 64 µL lysate.

#### Liquid Chromatography – Mass Spectrometry analysis

LC-MS analysis was performed using a Vanquish^™^ Flex^™^ Tandem UHPLC system (Thermo Fisher Scientific, UK) connected to a Q-Exactive^™^ Hybrid Quadrupole-Orbitrap^™^ Mass Spectrometer (Thermo Fisher Scientific, UK). Samples were injected at a volume of 5 µL and separated using one of two, ACQUITY Premier BEH C18 Columns with VanGuard FIT, 1.7 µm, 2.1 x 50 mm (Waters, 186009455) at a column temperature of 40°C and a flow rate of 700 µL min^-1^. Samples were analysed using a six-minute gradient method using water with 0.2% (v/v) formic acid (Optima^™^ LC/MS-grade, Thermo Fisher Scientific, UK) and acetonitrile (HPLC-grade, VWR) as the A and B mobile phases, respectively. Starting with 95% mobile phase A, 5% mobile phase B, the LC gradient was as follows: 0.0 →1.0 min (5% B), 1.0→2.0 min (5%→ 95% B), 2.0→5.0 min (95% B), 5.0→5.1 min (95%→5% B), 5.1→6.0 min (5% B). Flow from the column was sent to the mass spectrometer in a window set to between 0.5 and 4.5 minutes. Three consecutive injections of acetonitrile/water (80/20 (v/v)) were performed after each sample injection to prevent carry-over.

Mass spectrometry analysis was performed with electrospray ionisation in positive ionisation mode using the following source conditions: source gas flow = 55 arb. units; auxiliary gas flow = 10 arb. units; auxiliary gas heater; temperature = 310°C; spray voltage = 3.2 kV; capillary temperature = 320°C; S-lens RF level = 55.0 arb. units. Analysis was performed using a full scan MS method with 35,000 mass resolution at 200 *m/z*, and a scan window from 100-900 *m/z*. Target peaks were as follows: ATI-2307 *m/z* = 219.6468 [M + 2H]^2+^, Iminoctadine *m/z* = 178.6784 [M + 2H]^2+^, Pentamidine *m/z* = 171.1022 [M + 2H]^2+^.

Sample quantifications of ATI-2307, Iminoctadine and Pentamidine were performed using matrix-matched calibration curves with R^2^ values greater than 0.996. Calibration curves were prepared using serial dilutions of compound stock solutions in yeast lysate extracts. For Pentamidine and Iminoctadine, two calibration curves were prepared: one to quantify the wild-type samples, and one to quantify the *hol1*Δ samples. Calibration curves were calculated using Xcalibur^TM^ (Thermo Fisher Scientific, v4.6), and the *m/z* of the peaks for each calibration point were verified using Quan Browser software (Thermo Fisher Scientific, v4.6).

### Screening for separation-of-function mutations in *HOL1*

The CEN*/URA3* plasmid bearing *S. cerevisiae HOL1* (pBK958), a kind gift from Dr Thomas Dever (NIH, Bethesda, Maryland, USA), served as the template for PCR. The *ScHOL1* CDS and flanking UTR sequences were PCR-amplified using *Taq* DNA Polymerase, generating a product that contained 5’ and 3’ sequences homologous to the pBK958 backbone. *ScHOL1* CDS was excised from pBK958 using *Bsu36*I and *Age*I, and the resulting pBK958 backbone was co-transformed into the *spe1*Δ *hol1*Δ strain (ByK2037) with the *ScHOL1* PCR product. Transformants were selected on SC -URA 2% glucose, and subsequently replica plated onto SC -URA 2% ethanol medium containing 1 μM spermidine (Sigma-Aldrich, S2626) and 0.01 μg/mL ATI-2307. ATI-2307-resistant isolated colonies were then streaked onto SC -URA 2% ethanol + 1 μM spermidine + 0.01 μg/mL ATI-2307 plates for a second round of selection. Plasmids were isolated from ATI-2307-resistant colonies, sequenced, and re-transformed back into the *spe1*Δ *hol1*Δ strains prior to disc diffusion assays.

### Disc diffusion assays

Cells were grown to saturation in 5 mL SC -URA 2% ethanol 0.2% glucose medium supplemented with 1 mM spermidine and 100 nM pantothenic acid. 175 μL of each saturated culture was then plated on SC 2% ethanol supplemented with 1 μM spermidine. Discs (Thermo Fisher Scientific, 10066142) were wetted with 17 μL of each drug (ATI-2307 0.4 μg/mL; Pentamidine 80 μg/mL; Iminoctadine 30 μg/mL) and placed on the agar surface. Plates were incubated at 30°C for five days before imaging.

## Data availability

All data are available in the main text, supplementary figures, and in the genome browser links provided.

## Acknowledgements

The authors are grateful to Christian Coville-Cooke and Ying-Chen Lin for feedback on the manuscript and technical support; Anitha Nair and Yaan Jang for technical support; Jack Rice for technical support and assistance with LC-MS. M.T.K. was supported by an Industrial CASE (iCASE) Studentship, jointly funded by the BBSRC and Syngenta (grant number (BB/T008784/1). The Kornmann Lab is funded by Wellcome Trust grant 214291/Z/18/Z.

## Supplementary Figure Legends

**Figure S1.**
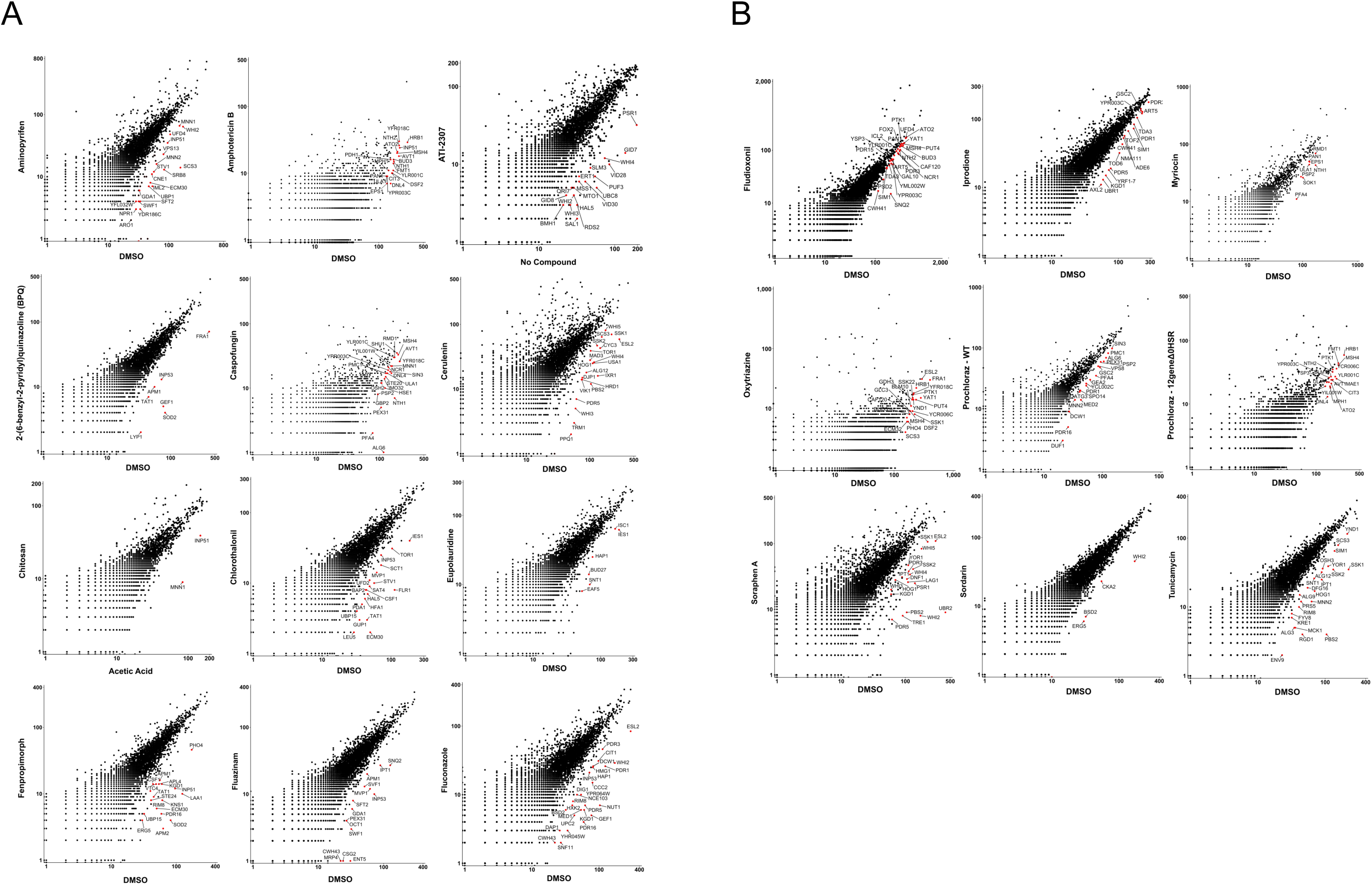
Scatter plots comparing the number of transposons mapping in the coding DNA sequence (CDS) of every gene in transposon libraries treated with each compound.

**Figure S2.**
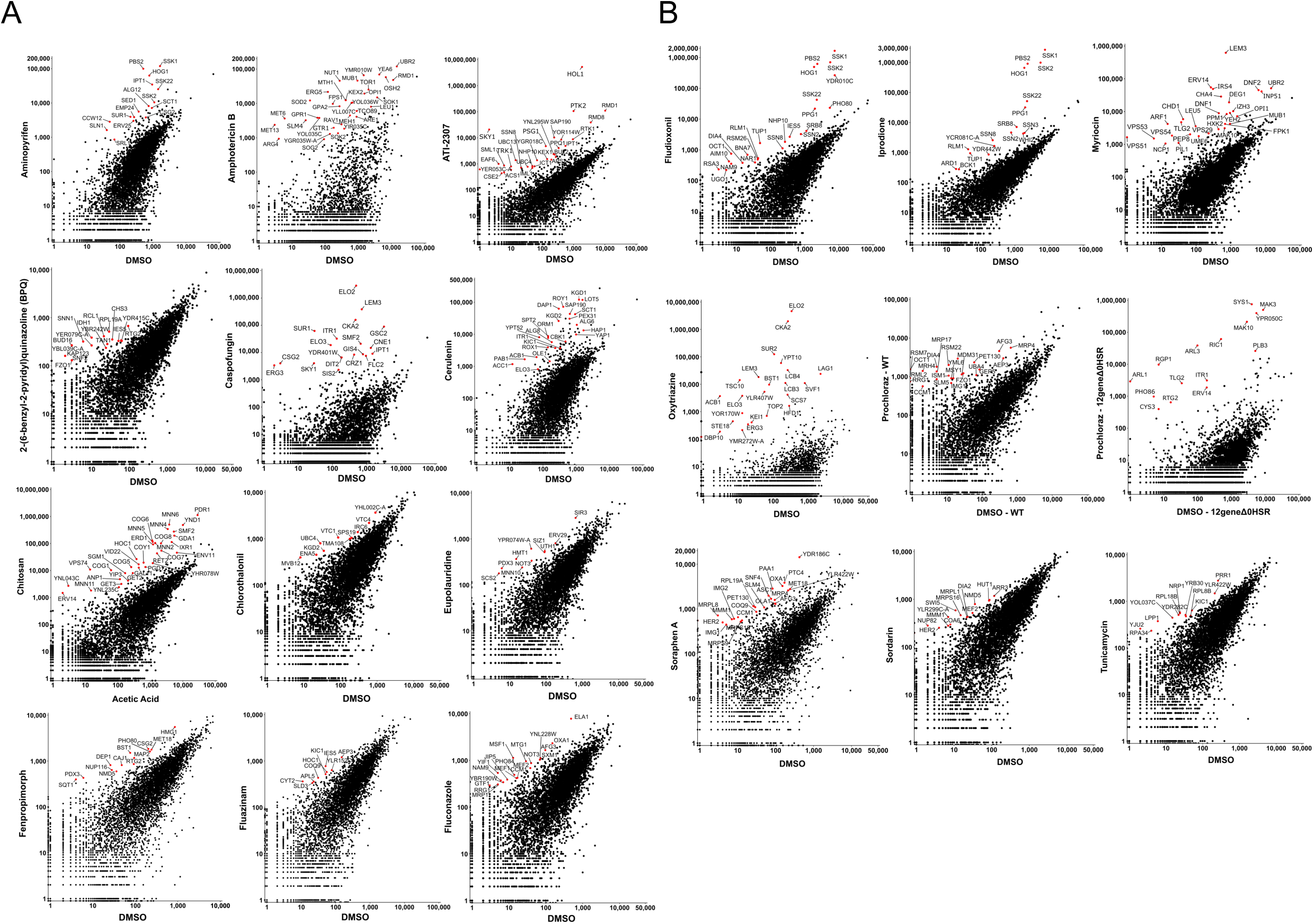
Scatter plots comparing the number of sequencing reads mapping in the coding DNA sequence (CDS) of every gene in transposon libraries treated with each compound.

**Figure S3.**
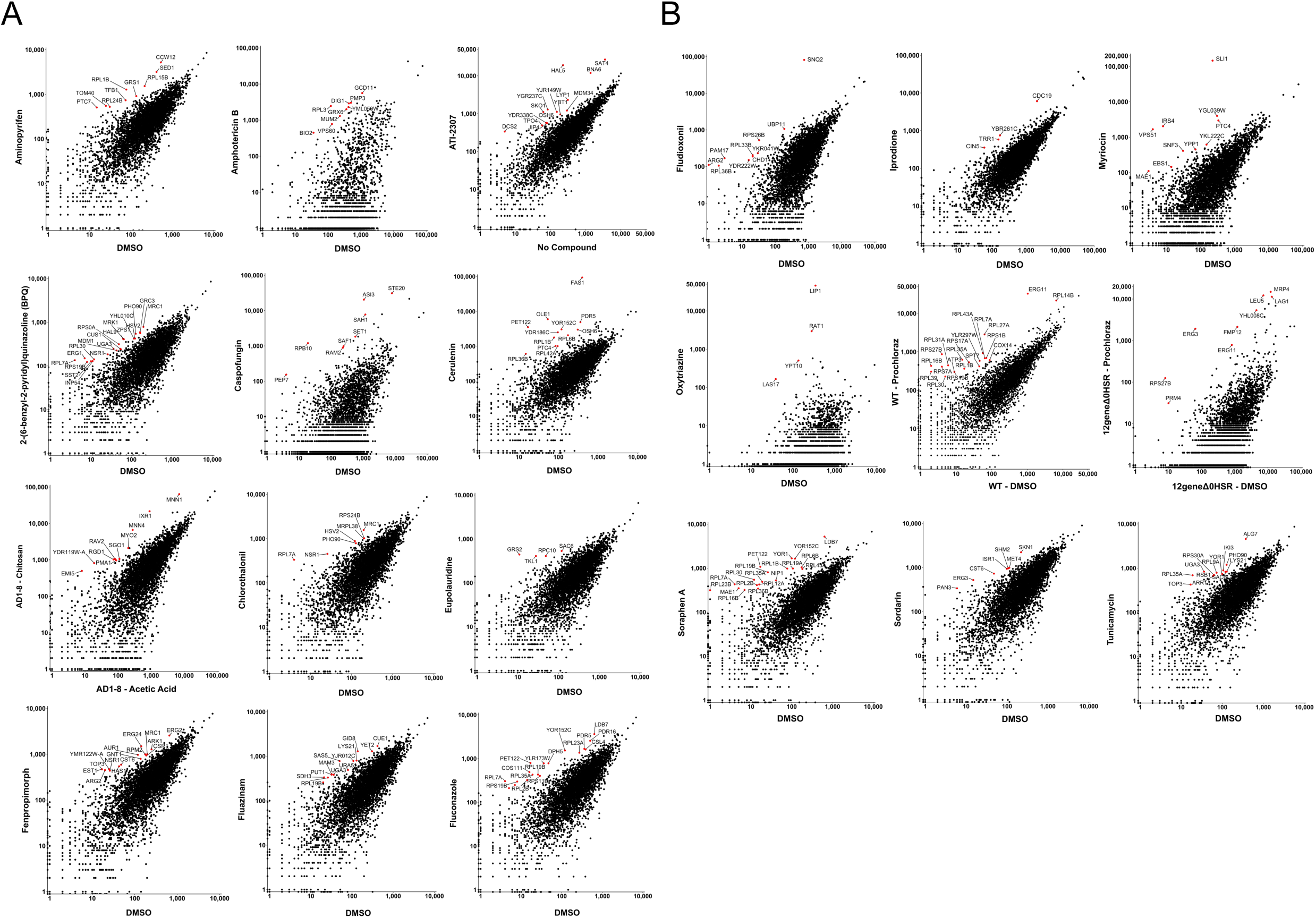
Scatter plots comparing the number of sequencing reads mapping to the promoters in transposon libraries treated with each compound. Promoters are defined as the 200 bp upstream of the transcription start sites, as assessed by Xu et al., (2009).

**Figure S4.**
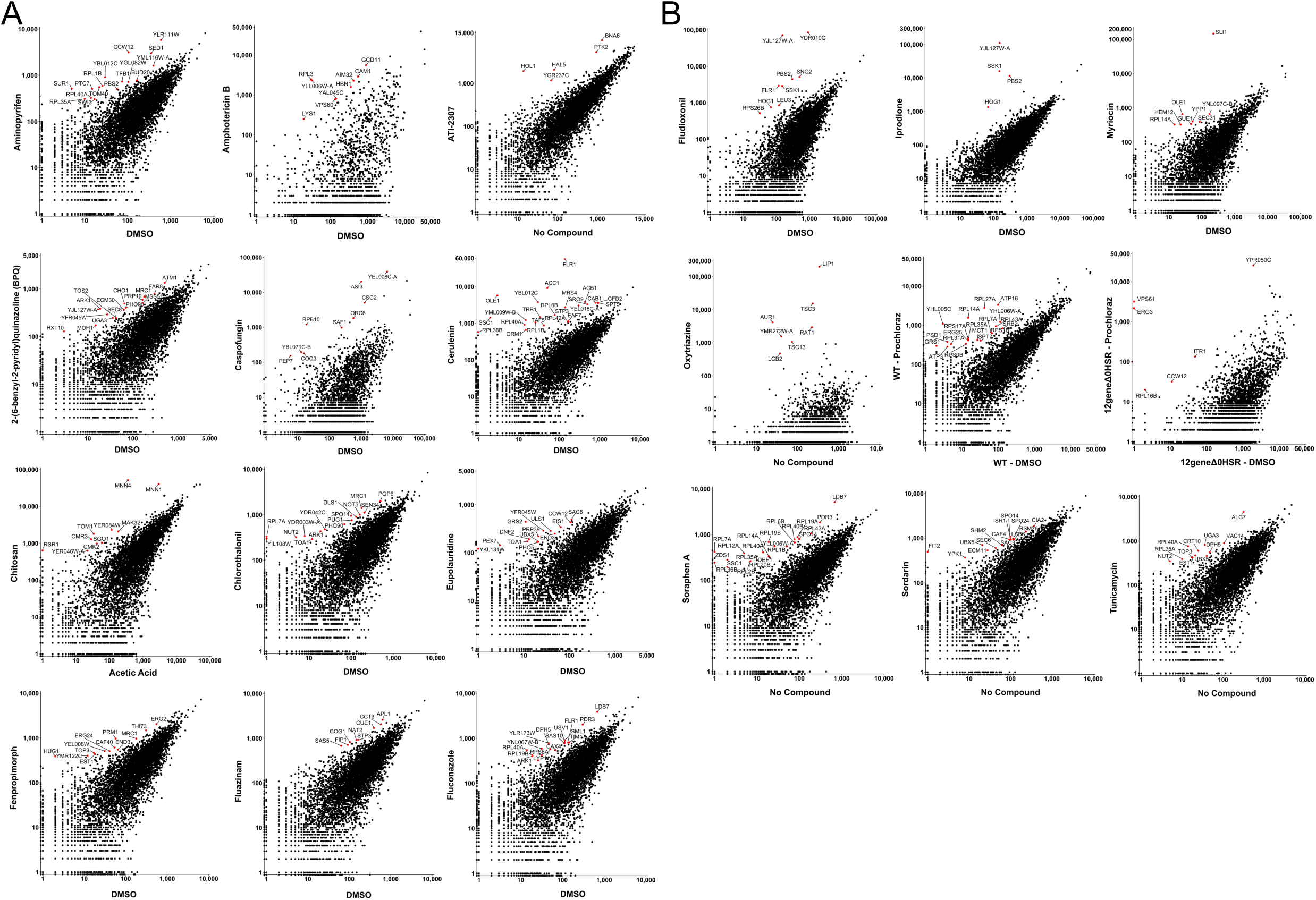
Scatter plots comparing the number of sequencing reads mapping to the promoters in transposon libraries treated with each compound. Promoters are defined as the 200 bp upstream of the translation start site.

**Figure S5.**
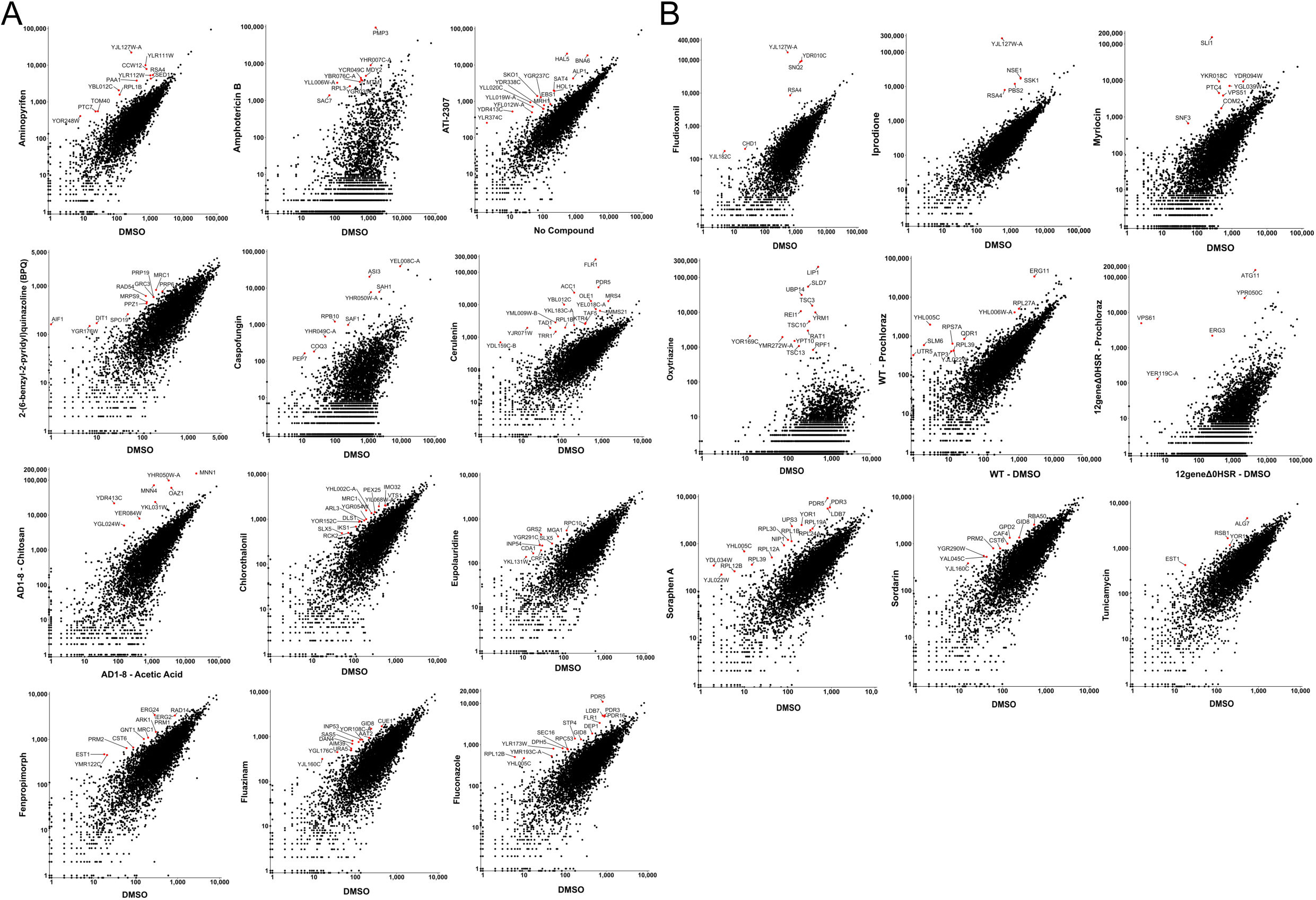
Scatter plots comparing the number of sequencing reads mapping to the promoters in transposon libraries treated with each compound. Promoters are defined as the 500 bp upstream of the translation start site.

**Figure S6.**
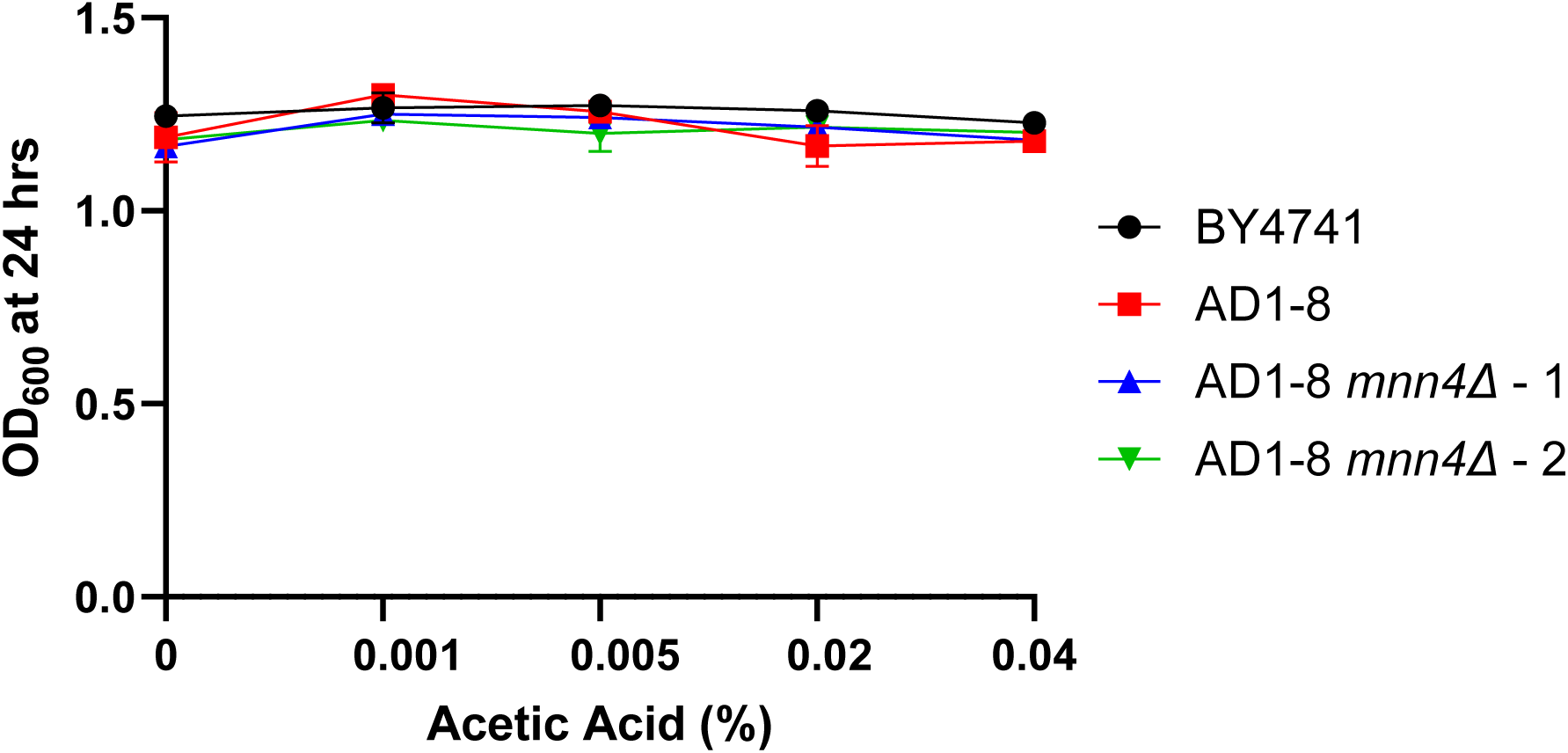
Growth inhibition curves of a BY4741 strain (ByK352), AD1-8, and two independent *mnn4*Δ AD1-8 strains treated with acetic acid in SC 2% glucose medium. Acetic acid concentrations (% v/v) correspond to those in each Chitosan treatment (i.e. the 200 μg/mL Chitosan treatment contained 0.04% acetic acid (v/v)). End-point OD_600_ measurements were recorded after 24 hours using a Bioscreen C™ instrument set at 30°C, continuous shaking. Each data point is the mean OD_600_ measurement for three technical replicates. Error bars show standard deviation (SD). Error bars for some measurements were too small to be displayed.

**Figure S7.**
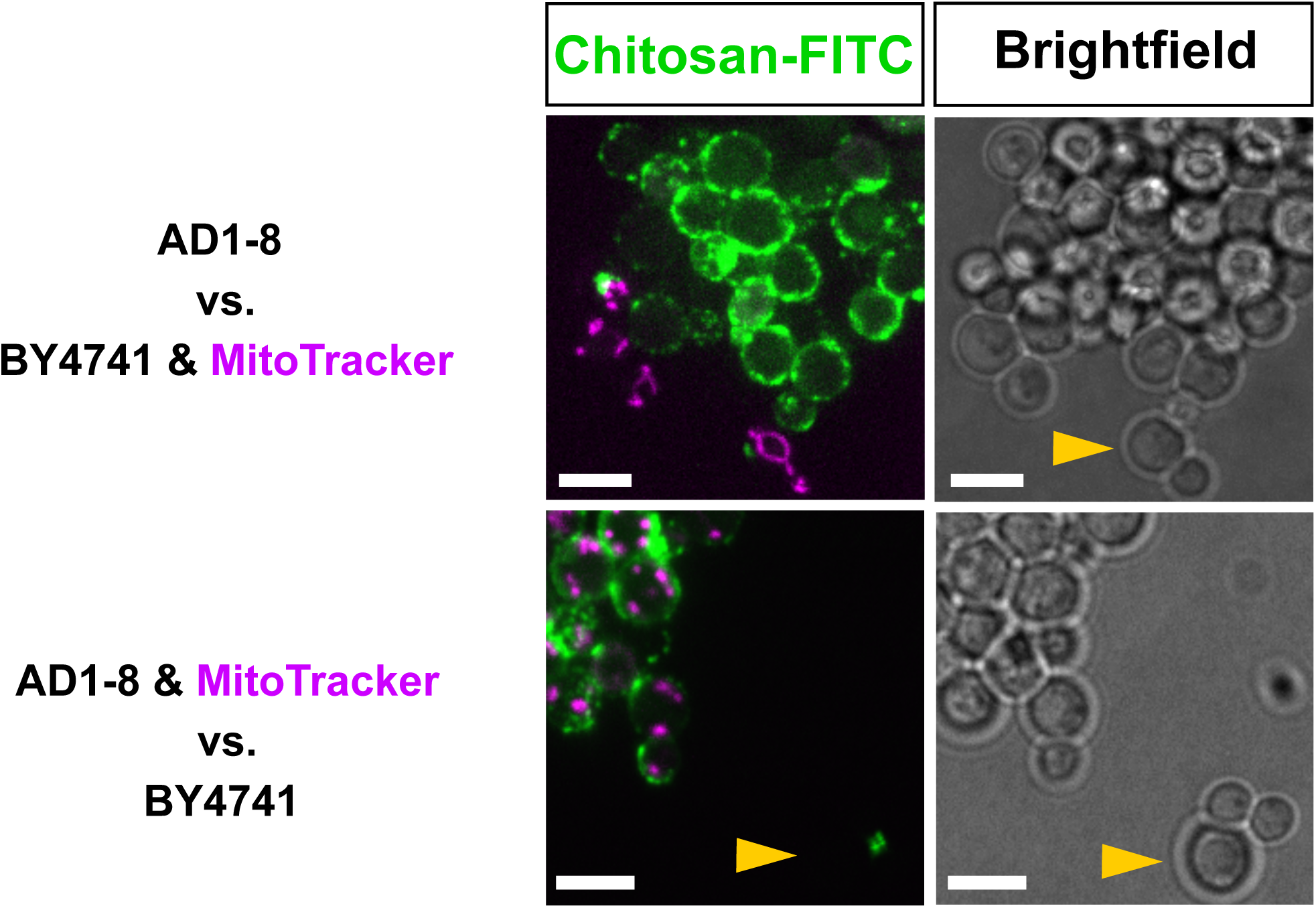
Representative images of AD1-8 and a BY4741 strain (ByK352) following treatment with 25 μg/mL Chitosan Fluorescein (green) for 2 hours in stationary phase in SC 2% glucose medium. MitoTracker (magenta) was applied at 0.1 μM. Yellow arrowheads indicate representative BY4741 cells. Images are Z-stacks. Scale bar represents 5 μm.

**Figure S8.**
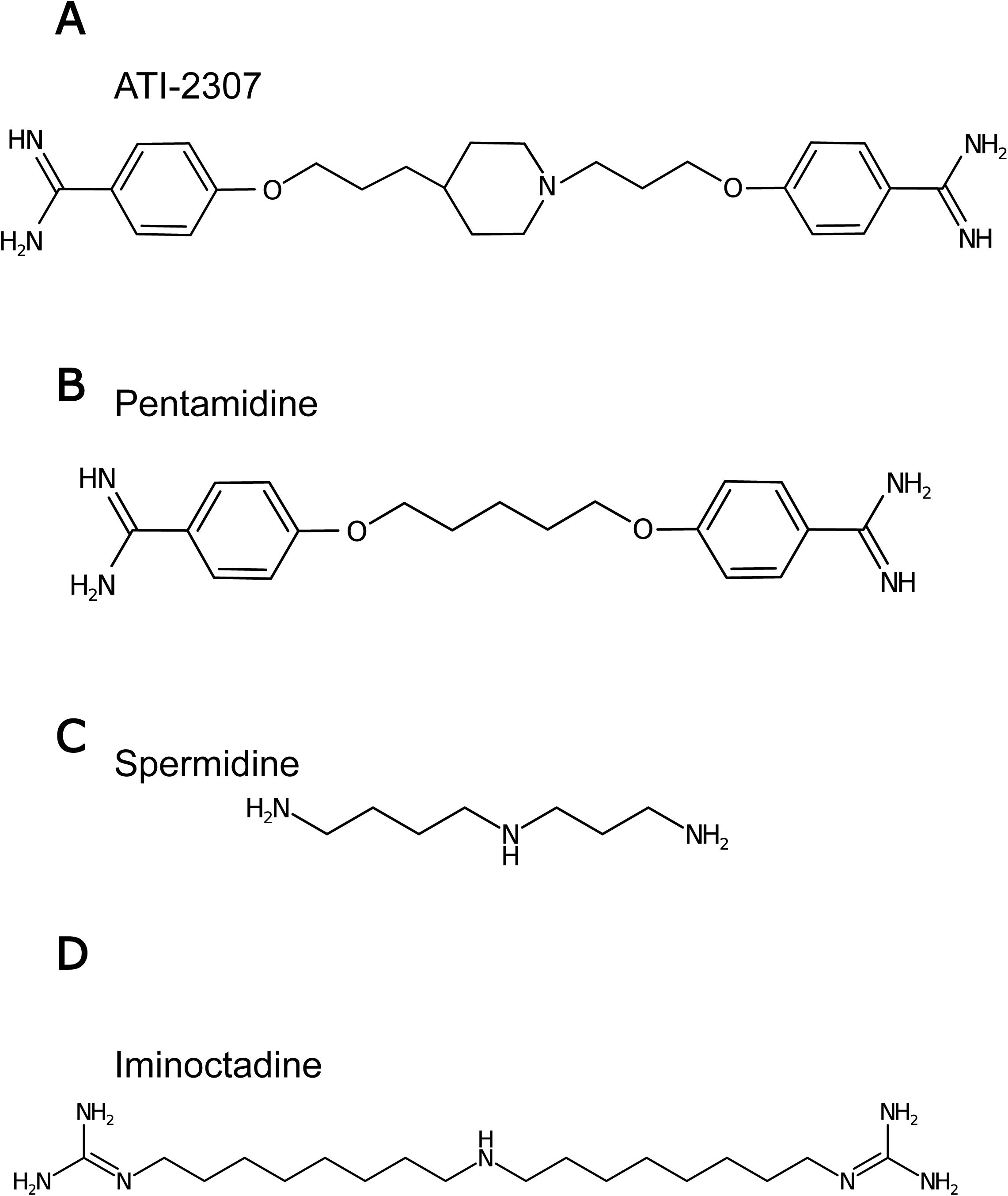
Chemical structures of A) ATI-2307 B) Pentamidine C) Spermidine D) Iminoctadine.

**Figure S9.**
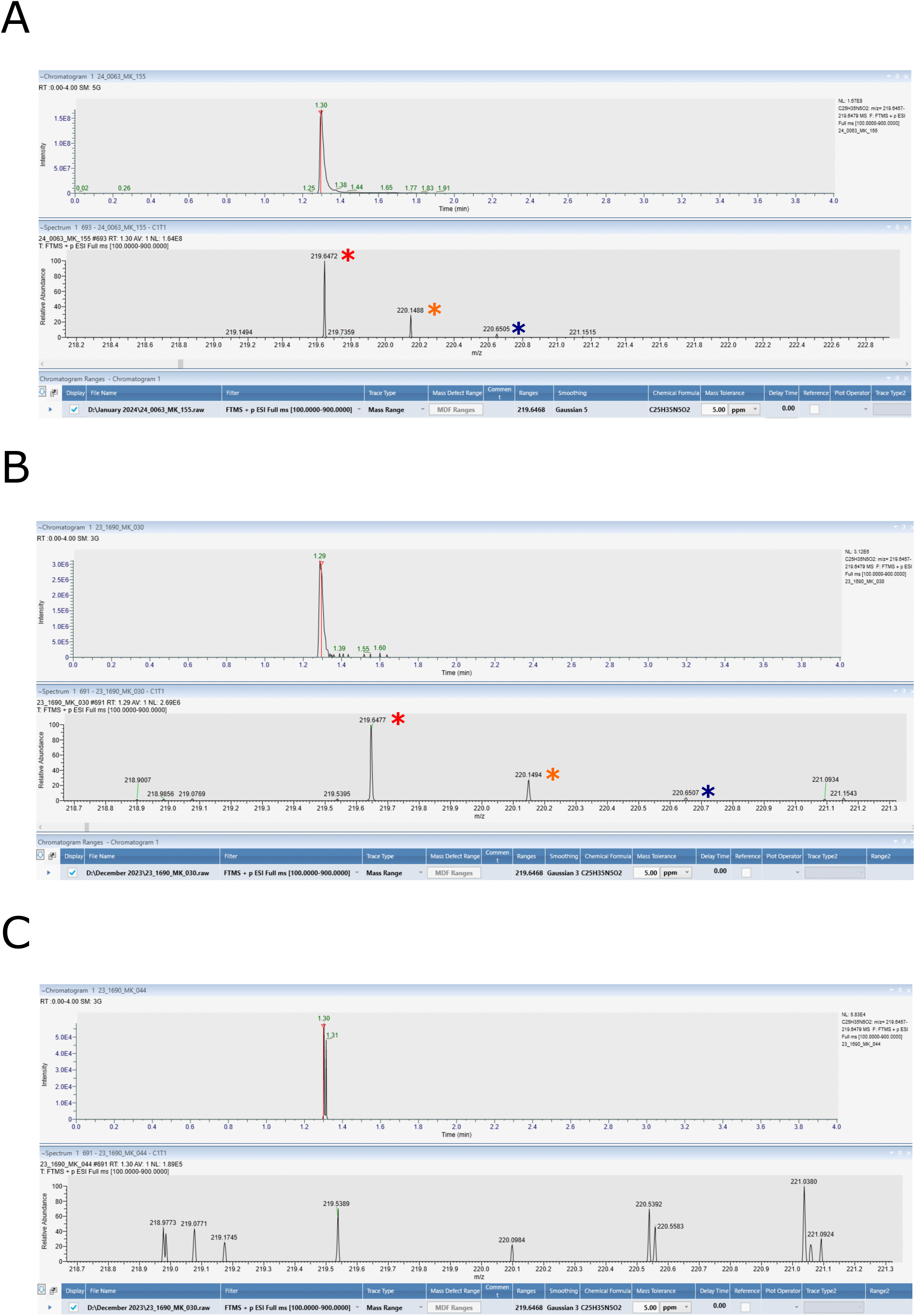
Chromatograms and mass spectra for A) yeast lysate spiked with 1.25. μg/mL ATI-2307 B) lysate obtained from ATI-2307-treated WT sample C) lysate obtained from ATI-2307-treated *hol1*Δ **sample**. ATI-2307 was identified with a retention time of 1.30-minutes. For mass spectrometry analysis, ATI-2307 was detected with a +2 charge. Red asterisks denote molecular ion peak ([M + 2H]^2+^) at m/z = 219.6472. Orange and dark blue asterisks denote isotope peaks. Note that ATI-2307 was not detected in the lysate from the *hol1*Δ sample.

**Figure S10.**
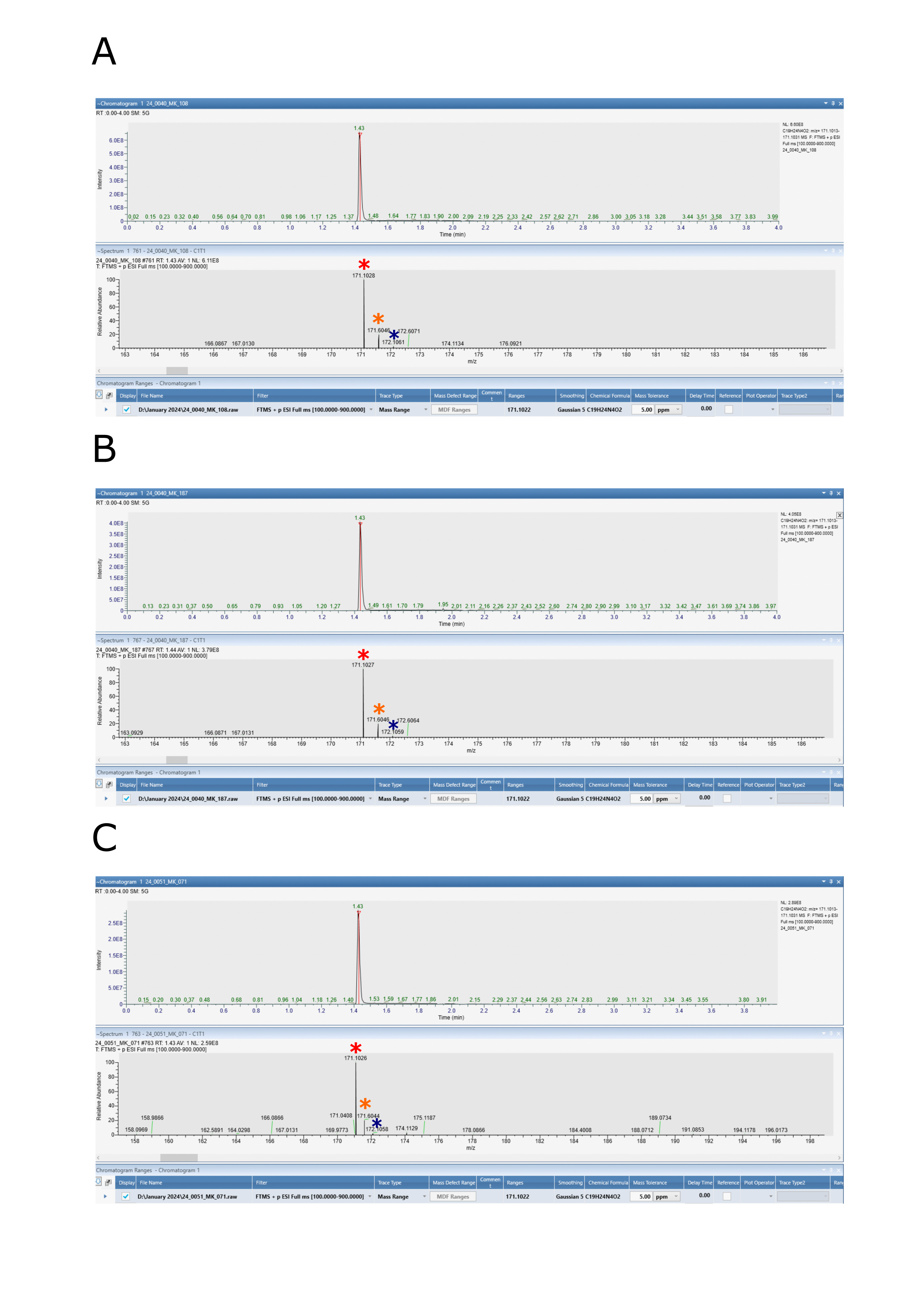
Chromatograms and mass spectra for A) yeast lysate spiked with 1.25. μg/mL Pentamidine B) lysate obtained from Pentamidine-treated WT sample (diluted 1/10 in yeast lysate) C) lysate obtained from Pentamidine-treated *hol1*Δ **sample**. Pentamidine was identified with a retention time of 1.43-minutes. For mass spectrometry analysis, Pentamidine was detected with a +2 charge. Red asterisks denote molecular ion peak ([M + 2H]^2+^) at m/z = 171.1027. Orange and dark blue asterisks denote isotope peaks.

**Figure S11.**
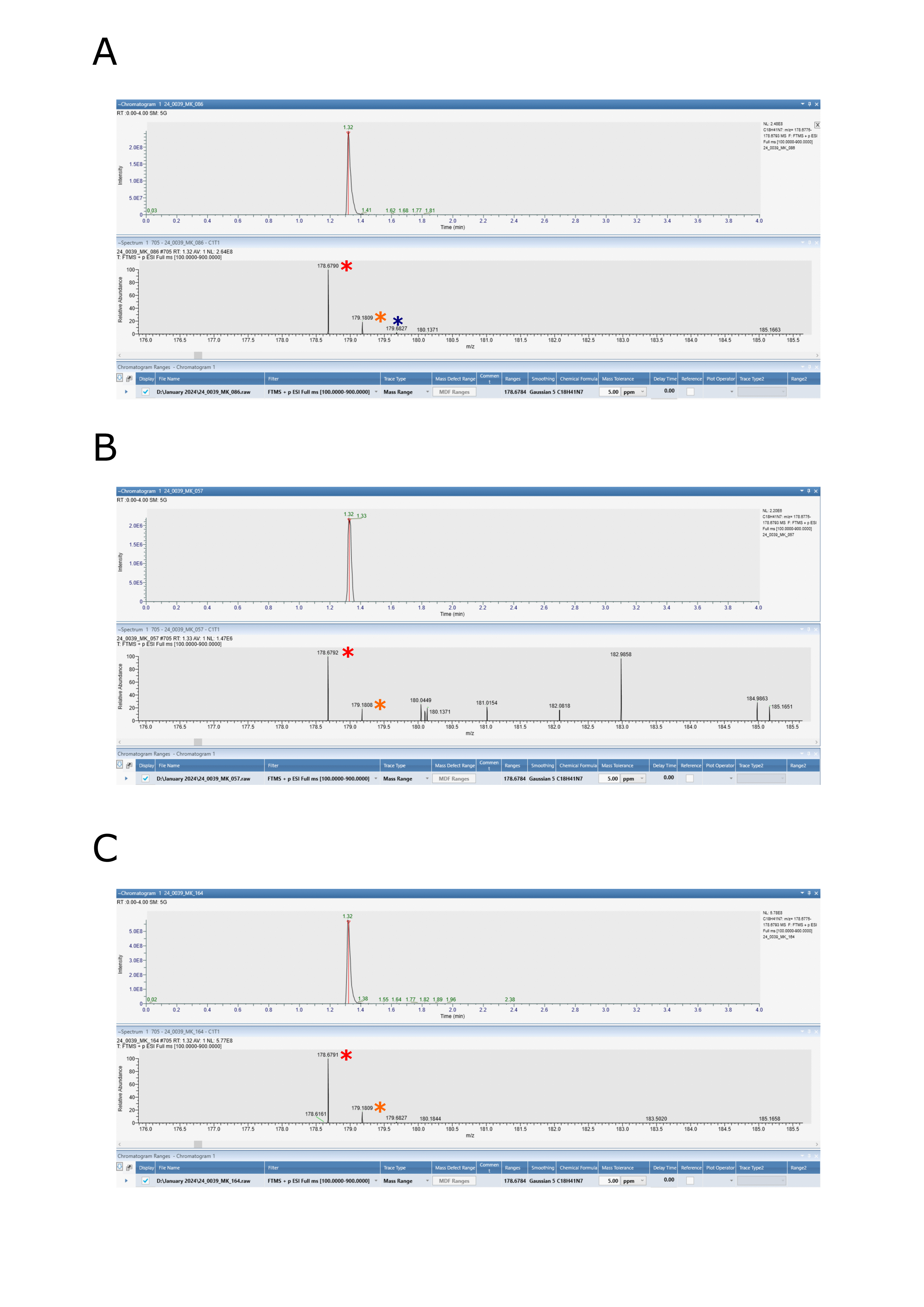
Chromatograms and mass spectra for A) yeast lysate spiked with 1.25. μg/mL Iminoctadine B) lysate obtained from Iminoctadine-treated WT sample (diluted 1/10 in yeast lysate) C) lysate obtained from Iminoctadine-treated *hol1*Δ **sample**. Iminoctadine was identified with a retention time of 1.32-minutes. For mass spectrometry analysis, Iminoctadine was detected with a +2 charge. Red asterisks denote molecular ion peak ([M + 2H]^2+^) at m/z = 178.6790. Orange and dark blue asterisks denote isotope peaks.

**Figure S12.**
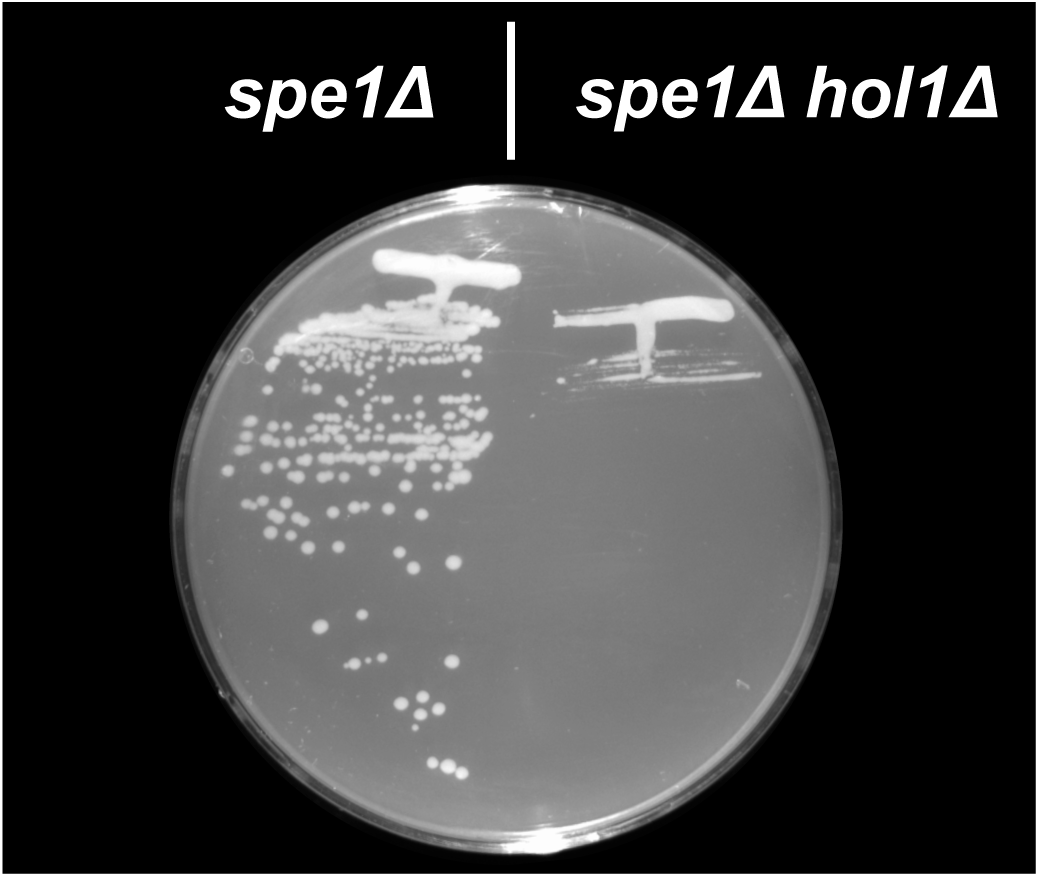
Streaks of *spe1*Δ and *spe1*Δ *hol1*Δ strains on SC 2% ethanol medium supplemented with 1 µM spermidine. Image taken after 6 days of incubation at 30 °C.

**Figure S13.**
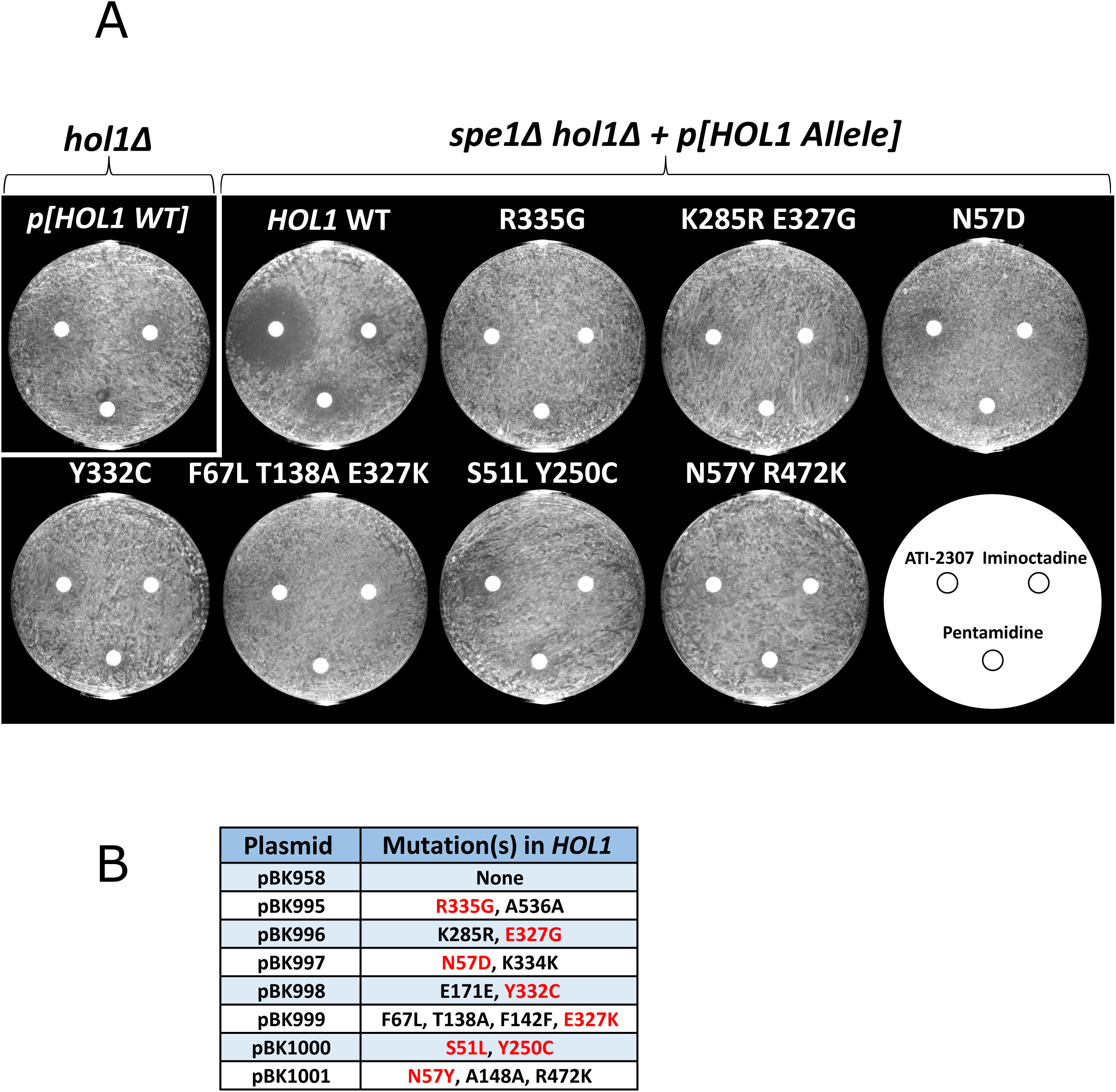
Disc diffusion assays testing ATI-2307, Pentamidine and Iminoctadine susceptibility of separation-of-function Hol1 mutants. **A)** *hol1*Δ transformed with pBK958 (*HOL1* WT) (top-left). *spe1*Δ *hol1*Δ transformed with pBK958 (*HOL1* WT) or one of seven plasmids (pBK995-pBK1001) encoding *HOL1* alleles bearing separation-of-function mutations. Non-synonymous mutations in *HOL1* for each plasmid are shown above the corresponding plate. For each transformant, 175 μL of saturated culture (OD_600_ ∼4) was plated on SC 2% ethanol medium supplemented with 1 μM spermidine. On each plate, discs were treated with 17 μL of 0.4 μg/mL ATI-2307 (top-left), 80 μg/mL Pentamidine (bottom) or 30 μg/mL Iminoctadine (top-right). Images taken after 5 days of incubation at 30°C. The *SPE1 hol1*Δ strain (top-left) did not require the plasmid-borne *HOL1* gene for viability on SC 2% ethanol + 1 μM spermidine medium. Hence, this strain can lose the plasmid and grow in the presence of the three drugs. In contrast, the *spe1*Δ strains required the plasmid-borne *HOL1* gene for growth on this medium. As such, the *spe1*Δ strain transformed with the WT *HOL1* gene was susceptible to the three drugs. **B)** Table of plasmids used in disc diffusion assay. Non-synonymous mutations shown in red were considered to be causal of the separation-of-function phenotype.

**Figure S14.**
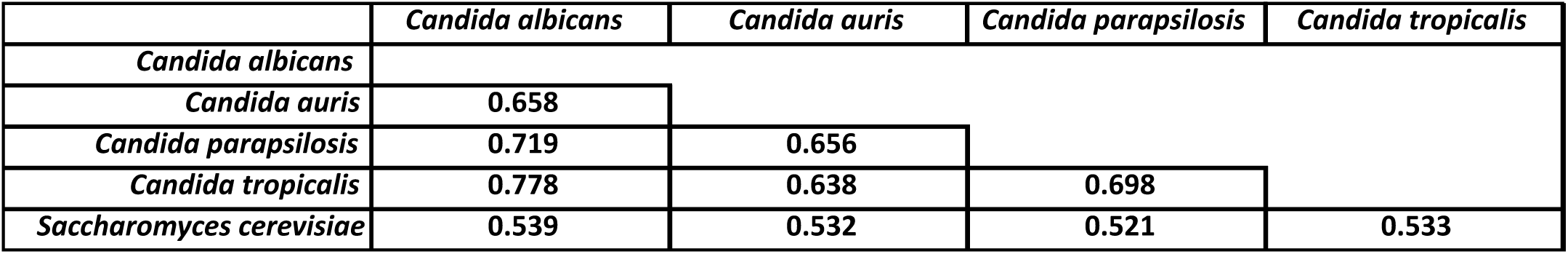
Pairwise sequence alignments of Hol1 protein sequences from *Candida* species and *S. cerevisiae*. Pairwise sequence alignments were performed using EMBOSS Needle. Hol1 protein sequences for each species were obtained from UniProt with the following identifiers: *C. albicans* Q5AP74; *C. auris* A0A890CZK8; *C. parapsilosis* G8BA41; *C. tropicalis* C5MB76; *S. cerevisiae* P53389.

