## Supplementary Methods for "SATAY-Based Chemogenomic Screening uncovers Antifungal Resistance Mechanisms and Key Determinants of ATI-2307 and Chitosan Sensitivity"

**Yeast media composition**

**Recipe A: 100x Amino Acid Mix (Department of Biochemistry, University of Oxford):**

L-Isoleucine (Sigma-Aldrich, W527602) 6 g/L, Adenine Hemisulfate salt (Sigma-Aldrich, A9126) 5.5 g/L, L-Arginine (Thermo Fisher Scientific, A15738) 2 g/L, L-Histidine monohydrochloride monohydrate (Sigma-Aldrich, H8000) 1 g/L, L-Leucine (Sigma-Aldrich, L8000) 6 g/L, L-Lysine monohydrochloride (Sigma-Aldrich, L8662) 4 g/L, L-Methionine (Sigma-Aldrich, M9625) 1 g/L, L-Phenylalanine (Sigma-Aldrich, P2126) 6 g/L, L-Threonine (Sigma-Aldrich, 89179) 5 g/L, L-Tryptophan (Sigma Aldrich, T0254) 4 g/L, L-Tyrosine (Sigma-Aldrich, 93829) 5.5 g/L, Uracil (Sigma-Aldrich, U1128) 5.5 g/L. Specific components were excluded for the preparation of drop-out media.

**Recipe B: 10x Amino Acid Mix (Institute for Biochemistry, ETH Zurich-like)*:**

L-Isoleucine (Sigma-Aldrich, W527602) 0.3 g/L, L-Valine (Sigma-Aldrich, V0500) 1.5 g/L, Adenine Hemisulfate salt (Sigma-Aldrich, A9126) 0.4 g/L, **L-Arginine (Thermo Fisher Scientific, A15738) 0.2 g/L**, **L-Histidine** **(Sigma-Aldrich, H8000) 0.2 g/L**, L-Leucine (Sigma-Aldrich, L8000) 1 g/L, L-Lysine monohydrochloride (Sigma-Aldrich, L8662) 0.3 g/L, L-Methionine (Sigma-Aldrich, M9625) 0.2 g/L, L-Phenylalanine (Sigma-Aldrich, P2126) 0.5 g/L, L-Threonine (Sigma-Aldrich, 89179) 2 g/L, L-Tryptophan (Sigma Aldrich, T0254) 0.4 g/L, L-Tyrosine (Sigma-Aldrich, 93829) 0.3 g/L, Uracil (Sigma-Aldrich, U1128) 0.2 g/L, **L-Glutamic Acid** (Sigma-Aldrich, G1251) 1 g/L, **L-Aspartic Acid** (Thermo Fisher Scientific, A13520) 1 g/L. Specific components were excluded for the preparation of drop-out media.

**Recipe C: 10x Amino Acid Mix (Institute for Biochemistry, ETH Zurich)*:**

L-Isoleucine (Sigma-Aldrich, I2752) 0.3 g/L, L-Valine (Carl Roth, 4879.3) 1.5 g/L, Adenine Hemisulfate salt (Sigma-Aldrich, A9126) 0.4 g/L, **L-Arginine monohydrochloride** **(Sigma-Aldrich, A5131) 0.2 g/L**, **L-Histidine monohydrochloride monohydrate** **(Sigma-Aldrich, H8125) 0.2 g/L**, L-Leucine (Sigma-Aldrich, L8000) 1 g/L, L-Lysine monohydrochloride (Sigma-Aldrich, L5626) 0.3 g/L, L-Methionine (Sigma-Aldrich, M9625) 0.2 g/L, L-Phenylalanine (Sigma-Aldrich, P2126) 0.5 g/L, L-Threonine (Sigma-Aldrich, T8625) 2 g/L, L-Tryptophan (Sigma-Aldrich, T0254) 0.4 g/L, L-Tyrosine (Sigma-Aldrich, T3754) 0.3 g/L, Uracil (Sigma-Aldrich, U0750) 0.2 g/L, **L-Glutamic Acid monosodium salt hydrate (Sigma-Aldrich, G1626) 1 g/L**, **L-Aspartic Acid sodium salt monohydrate** **(Sigma-Aldrich, 11195) 1 g/L**. Specific components were excluded for the preparation of drop-out media.

* Difference between recipe B and C shown in bold.

**SC medium (for recipe A):** 1x Amino Acid Mix (recipe A), 6.8 g/L Yeast Nitrogen Base without amino acids (BD Difco^TM^, 291920), carbon source (glucose, galactose, raffinose or ethanol) 2%.

**SC medium (for recipe B and C):** 1x Amino Acid Mix (recipe B or C), 1.7 g/L Yeast Nitrogen Base without ammonium sulfate without amino acids (BD Difco^TM^, 233520), 5 g/L ammonium sulfate (Sigma, A4418), carbon source (glucose, galactose, raffinose or ethanol) 2%.

**YPD medium:** Yeast Extract (Appleton Woods Ltd, DM832) 11 g/L, Peptone (BD Difco^TM^, 211677) 22 g/L, Adenine Hemisulfate salt (Sigma-Aldrich, A9126) 55 mg/L, Glucose 2%.

**YPA medium:** Yeast Extract (Appleton Woods Ltd, DM832) 10 g/L, Peptone (BD Difco^TM^, 211677) 20 g/L, Potassium acetate (Sigma-Aldrich, W292001) 10 g/L.

**Selective medium containing antibiotics:**

YPD-Geneticin: Geneticin (Fisher Scientific, 15444999) 0.2 mg/mL added after autoclaving

YPD-Nourseothricin: Nourseothricin (Melford, N51200-0.1) 0.1 mg/mL added after autoclaving

Solid medium was prepared using 2% Agar (BD Difco^TM^, 214030).
